# Motion-based depth estimation in *Drosophila*

**DOI:** 10.64898/2026.05.07.723493

**Authors:** Stefan Prech, Sophie Schmidt-Hamkens, Birte Zuidinga, Maria-Bianca Leonte, Lukas N. Groschner, Alexander Borst, Juergen Haag

## Abstract

Inferring three-dimensional structure from two-dimensional visual input requires a second perspective, either in time or in space. In flying animals, the first of these options lends itself as an ideal source for constructing internal representations of the environment, since their continuous self-motion automatically produces an optic flow. How neural circuits transform this signal into estimates of object distance, and what form such representations take, remains unknown. We address this problem in the visual system of the fruit fly. By immersing flies in a virtual environment while recording population-level calcium activity of motion-sensitive neurons, we extract an optic flow-based map of space. This map contains accurate estimates of object distance computed through spatial integration of elementary motion detector signals. Using electrophysiological recordings, we identify postsynaptic wide-field neurons as a cellular substrate for this operation. In behavioural experiments, we show that motion vision is essential for depth perception; without access to visual motion information, the free flight trajectories of motion-blind flies inevitably end in collisions. Our experiments link population-level neural activity to behaviourally relevant representations of environmental structure and demonstrate that motion vision is essential for navigation in three-dimensional space.

**One Sentence Summary:** Population activity of motion-sensitive neurons accurately captures object distance and is required for navigation in three dimensions.

## Main text

Eyesight is crucial for organisms that actively navigate their environment. The brain constructs, moment by moment, a mental map of its surroundings based on visual information. In the process of construction, it relies on visual features that hold implicit information about object distance: binocular disparity, relative size, occlusions, contrast, texture gradients, perspective, and motion parallax. The latter is of particular importance, because the vast majority of luminance changes that stimulate our retina come about not by objects moving in our surroundings, but by egomotion through predominantly static environments. When moving through space, nearby objects move faster across the retina than those at a distance, hence, creating heterogeneous patterns of visual motion^1–5^. As an airborne insect navigating complex three-dimensional spaces, *Drosophila melanogaster* is thought to rely primarily on these optic flow patterns to gauge distances, since alternative depth cues are either inaccessible or unreliable given its poor spatial resolution^6,7^.

Optic flow has two components: a rotational arising from turning about the body axis and a translational caused by motion through space along a straight trajectory. Much of the recent progress in understanding motion processing in the visual system of *Drosophila* comes from systematic experiments using uniform, rotational optic flow patterns^8–14^, which are independent of distance (Fig. 1a): When turning on the spot, the images of close and distant landmarks move across the retina at the same velocity. Translational flow patterns, by contrast, contain rich information about distance, depth, and *Umwelt* geometry^7,15,16^ (Fig. 1a). During egomotion, population activity of motion-sensitive neurons should, in theory, suffice to extract depth information^5,7,17^.

**Fig. 1.**
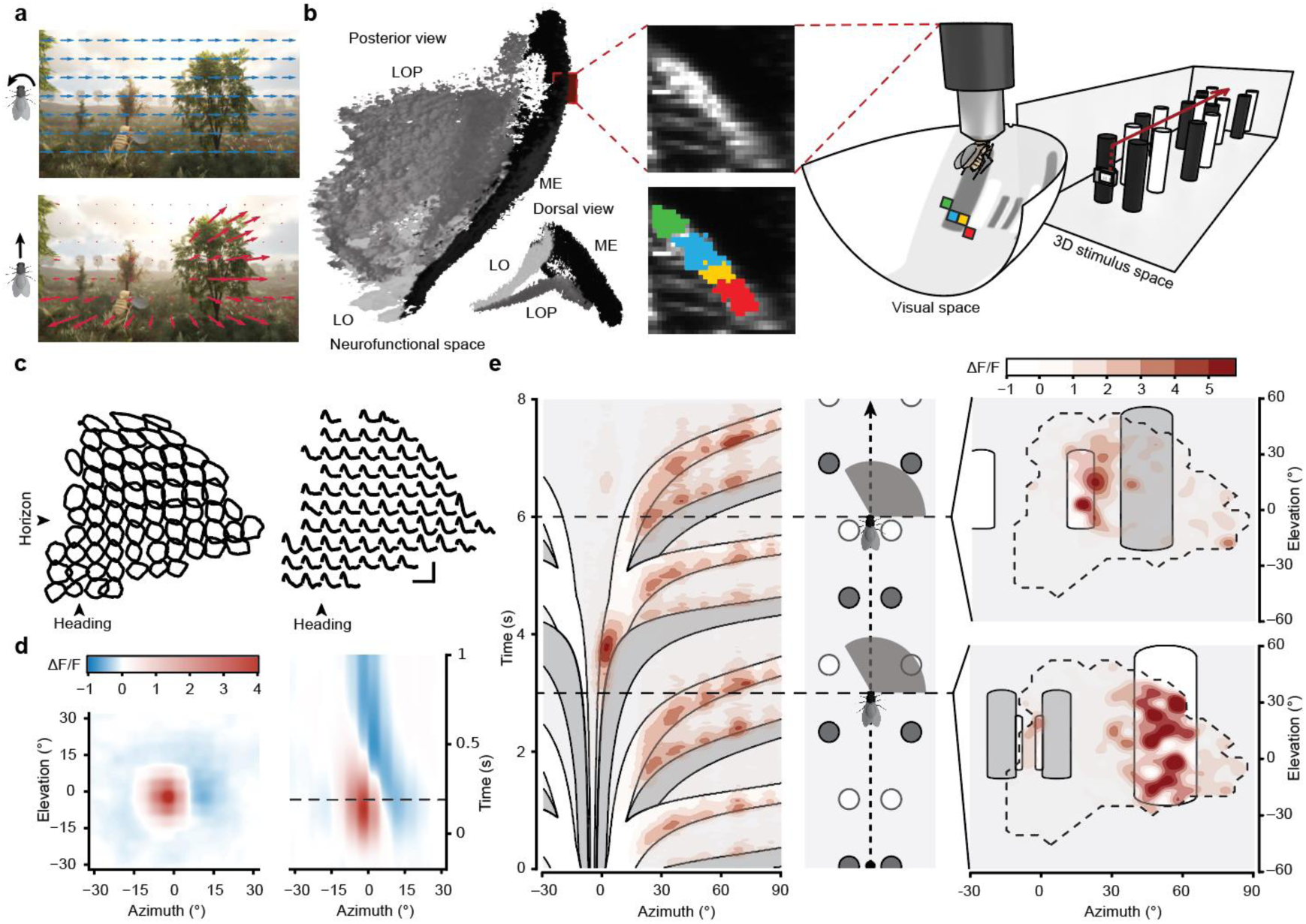
Reconstructing visual space from population activity of T4a neurons. **a**, During rotation (top), optic flow vectors (blue) are uniform across the visual field, reflecting distance-independent motion signals. During translation (bottom), flow vectors (red) vary in magnitude and direction depending on object distance. **b**, Functional–anatomical population imaging of T4a neurons. Left, two-photon calcium imaging of the right optic lobe during visual stimulation. ME, medulla; LO, lobula; LOP, lobula plate. T4a signals were sampled in the medulla. Middle, assignment of image pixels to virtual regions of interest (colours) based on spatial receptive field mapping using white-noise stimulation (see Methods). Right, schematic of the visual stimulus space and virtual flight trajectory. **c**, Distribution of receptive field properties across visual space. Left, receptive field full width at half maximum computed in 10° × 10° bins (total *n* = 1056 ROIs). Right, temporal impulse responses plotted as glyphs at corresponding locations. Horizontal scale bar, 10° ≙ 1.85 s; vertical scale bar, 10° ≙ 10 s.d. **d**, Averaged linear receptive fields. Left, spatial receptive field at 200 ms after stimulus onset, revealing an ON-centre with an asymmetric inhibitory surround. Right, spatio-temporal receptive field averaged across ±5° elevation. Horizontal dashed line, sampling time of the spatial receptive field on the left. **e**, Population responses during virtual passage through a scene with light and dark columns. Left, spatio-temporal activity plot (*ΔF/F*) and scene structure (greyscale; slope reflects stimulus velocity) averaged across ±25° elevation. Dashed lines, corresponding time points on the right. Centre, top view of the stimulus and fly trajectory at two example time points (3 s and 6 s). Right, reconstructed average activity maps (*n* = 12 flies) at corresponding time points, with stimulus geometry shown in greyscale and neural responses overlaid; dashed outlines indicate the reconstructed visual field.

Here, we test this hypothesis by viewing the world through the activity of motion-sensitive T4 and T5 neurons, the elementary motion detectors in the optic lobe of *Drosophila*. They respond in a direction-selective manner to moving luminance increments (ON, T4 cells) and decrements (OFF, T5 cells), respectively and segregate into four subtypes (a–d), each of which is sensitive to visual motion in one out of four cardinal directions^13^. Collectively, these neurons create a retinotopic array that tiles visual space. We map high-resolution population-wide calcium signals of T4a neurons sensitive to front-to-back motion onto visual space during translational optic flow in virtual visual environments. Our results demonstrate that the spatially integrated activity of motion detectors allows for the computation of object distance. Spatial integration takes place just one synapse downstream in wide-field neurons whose membrane potential dynamics contain distance-like information. An analysis of free-flight trajectories attests to the importance of motion vision for distance estimation: Motion-blind flies fail to steer clear of walls, and collide with obstacles at full speed.

### Reconstructing space from population-level activity of motion-sensitive neurons

We generated a GAL4 driver line that provided selective genetic access to T4a and T5a neurons (Extended Data Fig. 1a, b) and used it to target expression of the calcium indicator GCaMP8m to these front-to-back motion-sensitive neurons^13,18^. Genetic specificity combined with two-photon microscopy allowed us to record calcium signals from T4a dendrites in the medulla. Each experiment started with a binary white-noise stimulus displayed on a panoramic screen^19^ followed by a virtual flight trajectory through a 3D-rendered visual scene containing multiple objects at different distances (Fig. 1b). To accurately map the activity of densely interwoven T4a dendrites in the curved medulla onto visual space (Fig. 1b and Extended Data Fig. 2), we computed pixel-wise temporal cross-correlations between the white-noise stimulus (28 × 36 pixels, 5° per pixel) and the recorded fluorescence signals (32 × 32 pixels per plane). This produced a stimulus–response map linking each recorded pixel to its preferred stimulus location in visual space. Pixels with similar positions were grouped into virtual regions of interest, creating stable functional units for further analysis (Fig. 1b). By repeating this procedure across multiple image planes (*n* = 5–15 per fly) and flies (*n* = 12), we assembled a coherent and continuous map of neural activity aligned to a common visual coordinate system (Extended Data Figs. 1c, 2, 3 and 4).

Using this map, we examined how receptive field properties varied across visual space (Fig. 1c). Functional units were grouped into 10° × 10° bins, and receptive fields within each bin were registered and averaged. Receptive field full width at half maximum increased systematically from ∼8° in frontal regions to ∼12° peripherally, paralleling documented gradients in inter-ommatidial angle of the compound eye^20^. In contrast, temporal impulse responses were highly similar across space, indicating largely uniform temporal filtering (Fig. 1c).

The averaged spatio-temporal receptive field revealed an ON-centre embedded in a curved inhibitory structure, indicative of broad velocity tuning (Fig. 1d). Spatially, a broad but weak antagonistic surround coexisted with a strong localized inhibitory kernel, forming an asymmetric structure consistent with edge-aligned motion detection^21^. Temporally, the broad surround was short-lived, whereas the localized inhibitory kernel persisted and shifted toward the receptive field centre over time, consistent with directional filtering^10,22^.

Individual receptive fields were used to generate an anatomically grounded functional map to interpret neuronal responses during virtual passage of a scene containing black and white columns (Fig. 1b). In agreement with the selectivity of T4a neurons for front-to-back ON motion^13^, responses were only evoked by moving luminance increments, which occurred at the starting edges of white and at the ending edges of black columns (Fig. 1e and Supplementary video 1).

Response amplitudes were largely constant along the elevation, where the spatial arrangement of the scene remained unchanged, but varied substantially along the azimuth, where the luminance of the stimulus differed across positions (Fig. 1e). Along this axis, population activity was strongest, and had the largest spatial extent, for nearby objects. The responses were detectable over a wide range of speeds of up to 200°/s and at small spatial scales of as little as 5°/s, demonstrating the sensitivity of T4a neurons to motion across vastly different dynamics (Fig. 1e).

### Population responses capture object distance

The relationship between response amplitude and object distance (Fig. 1e) suggests that it is possible to infer distance based on T4a population activity. Testing this prediction, we presented flies with two stimuli that simulated translational motion towards a landmark. Each stimulus consisted of a white wall, that is, a single ON edge of identical luminance and size, ensuring full coverage of the field of view. The stimuli differed solely in the distance between the edge and the flight path and, hence, in the optic flow pattern on the retina. We chose a single edge because it represents the most reductionist discrete motion cue, allowing distance-dependent motion structure to be isolated while eliminating confounds from texture, contrast, or object identity.

To visualize how population activity evolved over the course of an approach, we superimposed reconstructed spatial activity profiles onto the visual scene at different time points (Fig. 2a). Both the amplitude and the spatial extent of the population response grew monotonically with decreasing distance up until the fly-by point, when the spatial extent reached 40° in azimuth. Along the elevation axis, activation remained evenly distributed throughout the approach, consistent with the uniform vertical expanse of the edge.

**Figure 2.**
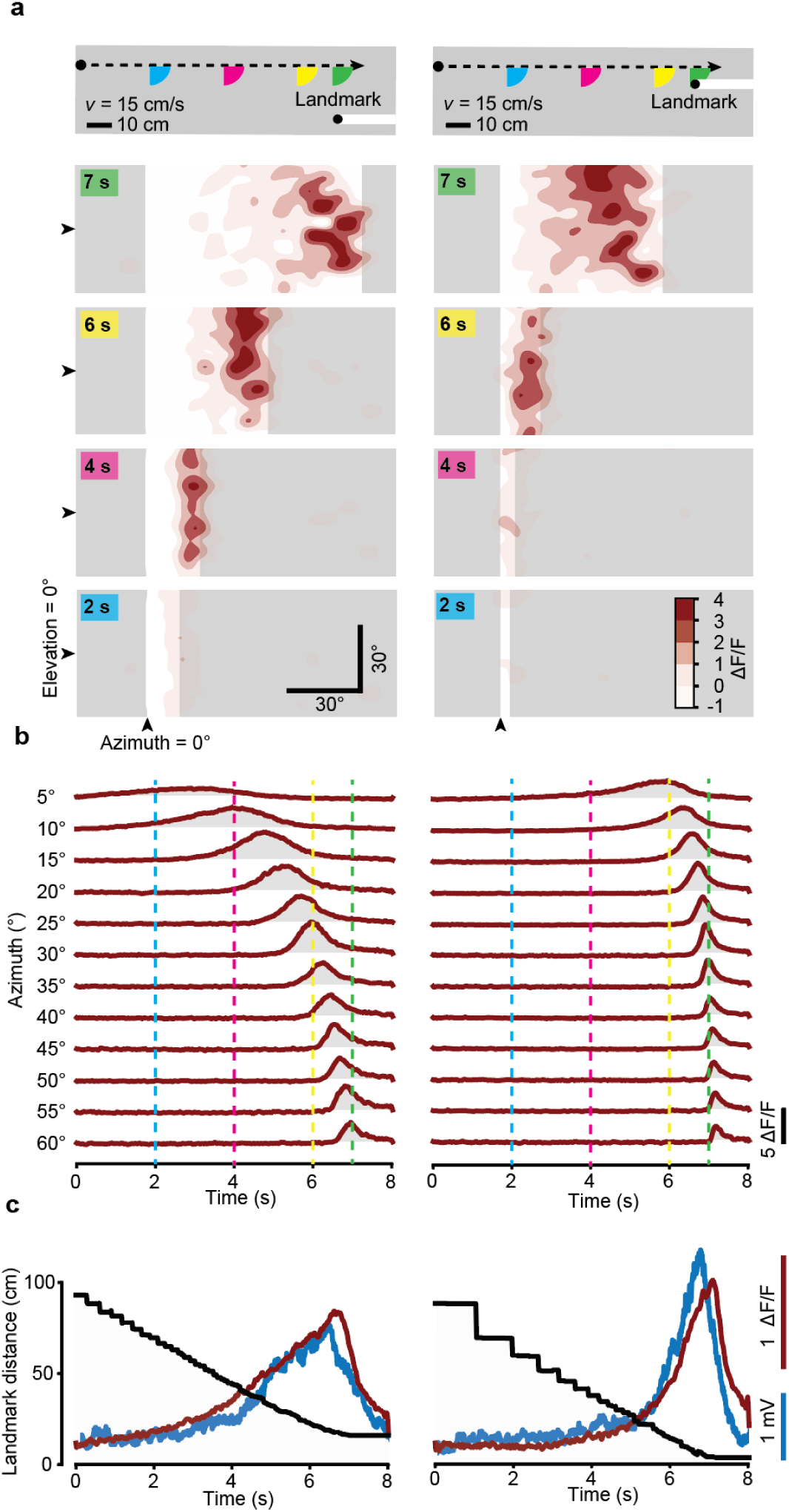
Population responses encode landmark distance during translational motion. **a**, Top, flies were sent on virtual trajectories (dashed lines) with a constant velocity (*v*) of 15 cm/s towards a landmark with a large (20 cm; left) and a small spatial offset (5 cm; right). Coloured angles, views at the respective timepoints. Bottom, T4a population calcium activity mapped onto visual space at different time points during simulated translations. Heatmaps show calcium activity (Δ*F*/*F*) in visual coordinates at 2, 4, 6, and 7 s during the approach. Colours correspond to time points indicated on top. **b**, T4a calcium signals (Δ*F*/*F*) at different azimuthal positions during the virtual fly-by at a large (left) and a small spatial offset (right). Azimuthal positions denote the centres of 5° bins, spaced at 5° intervals. Dashed vertical lines, time points shown in **a**. **c**, Spatially integrated population calcium responses of T4a neurons (red) membrane potential of postsynaptic HSE cells (blue; *n* = 10) and Euclidean distance between the fly and the landmark (black) during virtual translation.

Spatio-temporal response maps revealed a coherent wave of activity progressing along the azimuth in synchrony with the edge (Fig. 2b). Small retinal image shifts, consistent with a distant landmark, left activity confined to the position of the edge. Larger displacements, consistent with a nearby landmark, in contrast, resulted in simultaneous activation of multiple neighbouring azimuthal positions and temporal compression of responses (Fig. 2a, b).

Given the geometry-dependent evolution of activity across azimuthal positions, we next asked whether spatial integration of these responses could provide an accurate estimate of landmark proximity. Summing calcium responses over a range of positions from 0 to 75° in azimuth and ±25° in elevation, revealed a monotonic increase in activity as the landmark approached (Fig. 2c). This increase reflected the cumulative recruitment of spatially distributed elementary motion detectors as the edge traversed the visual field. The area used for spatial integration approximated the wide-field pooling performed by downstream neurons such as the horizontal system equatorial (HSE) cell^23^. To test if the HSE cell indeed carries a signal tied to landmark proximity, we used patch-clamp experiments to record membrane potential changes of HSE cells in response to the virtual translation past the landmark at different distances. Both pooled T4a calcium responses and HSE membrane potential dynamics were negatively correlated with the Euclidean distance to the landmark across conditions (*r*T4a = −0.93 and −0.68 ; *r*HSE = −0.78 and −0.75 for high and low offset, respectively; Fig. 2c). This indicates that spatial integration of local motion signals can transform distributed population activity into a distance-sensitive metric.

### Spatial integration supports accurate estimates of distance at close range

While spatially integrated T4a signals are correlated with landmark distance, it remains unclear whether distance information is already present at the level of individual detectors or emerges only through population-level interactions. To test this, we built a model (see Methods) consisting of an array of elementary motion detectors with T4-like dynamics (Fig. 3a–c). The model was fed with a simulated vertical edge travelling across the visual field. The mean Euclidean distance between the observer and the edge was varied systematically, while the lateral motion trajectory remained constant. Small retinal displacements elicited graded responses in individual motion detectors, whereas larger displacements, consistent with close objects, resulted in local response saturation (Fig. 3b). Spatial integration over multiple motion detectors yielded a monotonic increase in activity with decreasing distance. This relationship was robust to changes in motion speed (Fig. 3c). In the model, just like in calcium imaging experiments, proximity information was only weakly represented in the response amplitudes of single detectors due to rapid saturation, but emerged through the progressive recruitment of neighbouring detectors as distance decreased.

**Fig. 3.**
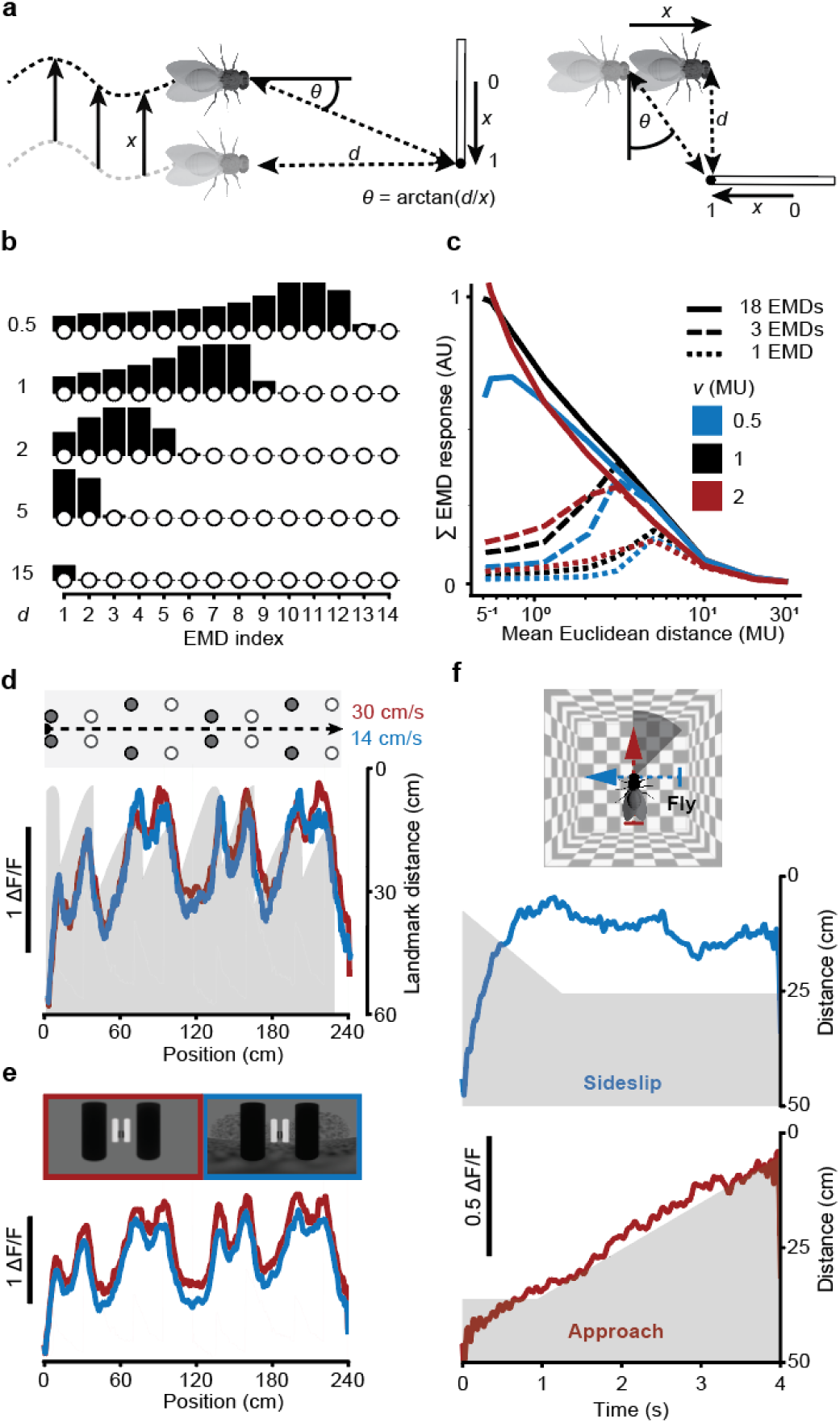
Robust encoding of landmark proximity through spatial integration of motion detector signals. **a**, Geometric configuration of the stimulus. A vertical edge at distance *d* and lateral offset *x* translates horizontally, generating a viewing angle *θ* = arctan(*d*/*x*). Comparable angular motion can arise either from small sideslip relative to a distant object (*d* ≫ *x*) or from forward translation towards nearby structure (*x* ≳ *d*). This geometric ambiguity necessitates encoding across a broad distance range, as both conditions may occur during natural flight. **b**, Individual elementary motion detector (EMD) responses at the end of a constant translation *x*: 0→1 with a velocity *v* = 1 (model units) for different distances (*d*). Bars, activity at time point *tend*; circles, EMDs. Decreasing distance recruited additional neighbouring detectors, while single-detector responses saturated. **c**, Linearly summed responses across 1, 3, or 18 detectors averaged at time point *tend* and plotted against mean Euclidean distance (log₁₀ scale). Spatial integration yielded a monotonic increase in population activity with decreasing distance, which was stable over a range of velocities (*v* = 0.5, 1, 2). AU, arbitrary units; MU, model units; EMD, elementary motion detector. **d**, Mean integrated T4a calcium responses (Δ*F*/*F*; *n* = 12 flies) during passage of a multi-landmark scene at low (blue) and high velocity (red). Activity was summed over 0–75° in azimuth and ±40° in elevation. Grey shading indicates distance to the nearest ON edge within the respective window. Responses increased as landmark distance decreased at both velocities. **e**, Responses during passage through the same scene as in **d** presented with low (red) and high (blue) background contrast (mean, *n* = 9). The addition of high-contrast noise to walls and ground did not affect response amplitudes. **f**, Sideslip and approach were simulated in a virtual checkerboard environment (top). Mean responses (*ΔF/F*, *n* = 12) were integrated over the same field of view used to compute minimum wall distance (grey; ray-based depth; 0–45° in azimuth and ±40° in elevation). During sideslip (centre), distance remained stable. During forward translation (bottom), distance and visible wall area decreased.

### Integrated responses track landmark proximity across velocities and scenes

To test whether distance-like signals obtained by spatial integration generalise to different velocities and visual scene statistics, we examined population responses to translational motion at varying speeds and scene contrast (Fig. 3d–f,).

We first revisited the stimulus used in Fig. 1, which contained multiple pillars at varying distances and azimuthal positions, including both bright and dark edges and partial occlusions (Fig. 3d). T4a population activity was integrated across an azimuthal range of 0–75°, analogous to the integration performed in Fig. 2c, and compared to the Euclidean distance of the nearest ON edges in the stimulus. For both velocities tested, the integrated neural response increased as the distance to an ON edge decreased (Fig. 3d). This relationship was consistent across all edges in the scene, indicating that the integrated signal robustly reflected landmark proximity. For the two velocities, the response profiles were highly similar in shape, suggesting that the integrated response was insensitive to changes in translational speed over the tested range (*r* = 0.95; ±0.1 *ΔF/F*; Two One-Sided Tests, *P* < 0.05).

To assess the influence of local contrast on this distance-like signal, we compared responses to the same complex scene stimulus presented either with or without additional background contrast (Fig. 3e).

Integrated responses in the high-contrast condition exhibited a significant reduction in amplitude compared to the low-contrast condition (paired t-test, *P* < 0.001). This difference exceeded predefined equivalence bounds (±0.1 *ΔF/F*; Two One-Sided Tests, *P* > 0.05). Despite this global signal attenuation, the temporal profile and distance dependence of the integrated response remained unchanged (*r* = 0.98; Fig. 3e). Thus, while strong background contrast modulated response gain, the geometric relationship between integrated activity and landmark distance was preserved under different contrast conditions.

Next, we asked whether the integrated T4a response primarily reflects object distance or instead scales with the number of visible landmarks within the integration region. To address this question, we placed the fly in a virtual box with checkerboard-patterned walls and compared two translational motion conditions at constant speed (Fig. 3f). During sideslip, both the distance to the wall facing the fly and the amount of visible structure within the integration region remained approximately constant. In contrast, during forward motion toward a wall, distance decreased approximately linearly over time, while the amount of visible structure progressively declined as a consequence of projective geometry. Despite this reduction in visible structure, neural responses increased smoothly during forward motion (Fig. 3f), indicating that the integrated signal does not simply scale with the number of visible landmarks.

### Natural flight-replay validates population-based distance encoding

Having established that spatially integrated motion responses provide a robust readout of landmark proximity across geometries and velocities, we next sought to test whether this signal persists under naturalistic conditions. To approximate natural flight as closely as possible, we used a representative three-dimensional flight trajectory of an unrestrained wild-caught fly inside a box with checkerboard-patterned walls^24^. The trajectory was selected from the longest recorded flights based on vertical stability, target height consistency, locomotor characteristics, and variation in scene depth (Extended Data Fig. 5a–c). We created a digital twin of the enclosure and rendered, frame by frame, the scene experienced by the fly, which served as visual stimulus during neural recordings. The flight kinematics featured characteristic straight flight segments interspersed with rapid saccadic turns (Fig. 4a), accompanied by substantial variations in flight speed (Extended Data Fig. 5d, e). This approach provided precise kinematic information and a pixel-wise representation of scene depth for every frame of the visual stimulus. The reconstructed trajectory was segmented into discrete, well-defined inter-saccadic flight segments during which the fly approached a wall monotonically (Fig. 4a). For one exemplary approach phase, we overlaid the reconstructed T4a population activity onto the corresponding visual scene at three time points (Fig. 4b and Supplementary video 2). Neural calcium responses increased progressively as the fly approached the wall. The increase was spatially selective and driven by regions exhibiting a decrease in distance, while regions without substantial distance change showed little modulation in response amplitude.

**Fig. 4.**
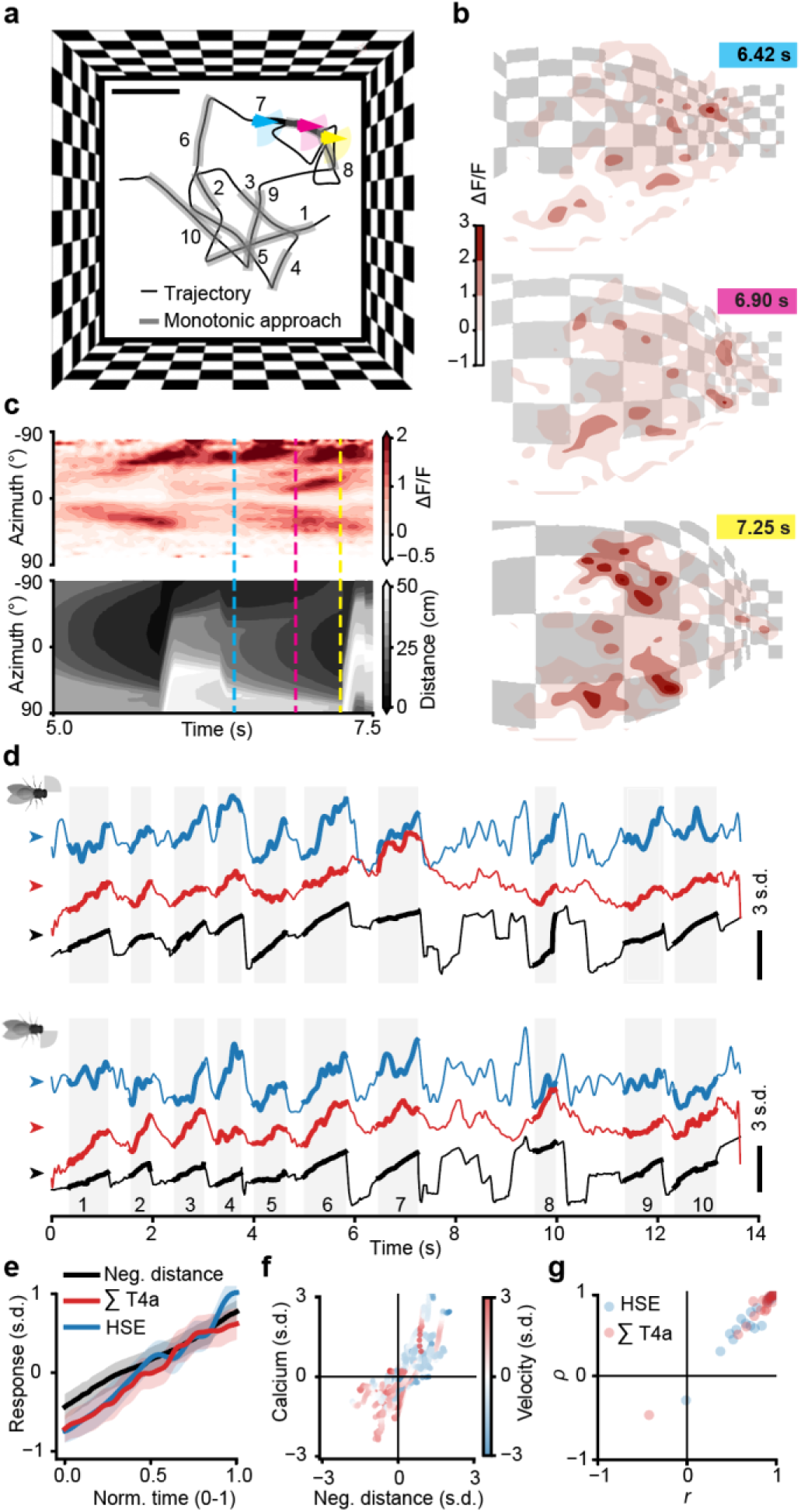
Population-based encoding of distance during naturalistic flight replay. **a**, Top view of the square arena used for free flight path reconstruction (black). Grey segments indicate inter-saccadic flight phases during which the fly approached a wall monotonically. Coloured arrows mark three time points within one example approach episode for analysis of neural responses in **b**. Numbers denote consecutive approach segments. Scale bar, 10 cm. **b**, Visual scenes experienced by the fly (grey) and reconstructed population activity of T4a neurons (red) at the time points indicated in **a** **c**, Spatiotemporal structure of reconstructed T4a population responses across both eyes as a function of time and azimuth averaged over elevations of ±30° (top) and wall distance obtained from a horizontal slice of the reconstructed depth map (bottom) during wall approaches 6 and 7 in **a**. Vertical dashed lines, time points indicated in **a**. **d**, HSE membrane potential (blue), spatially integrated T4a calcium signals (red), and negative wall distance (black) as *z*-scores during virtual transition of the flight path in **a** for the left (top) and right eye (bottom). Grey shading, monotonic approach phases; arrowheads, 0 s.d. **e**, HSE membrane potential (blue, *n* = 9 flies), population calcium signal (red, *n* = 9 flies) and negative stimulus distance (black) during monotonic wall approaches averaged across approaches (*n* = 20; original and mirrored stimuli). Time was normalized to approach duration (0–1). Data are mean ± s.e.m. **f**, *Z*-scored population calcium activity as a function of negative distance across monotonic approach phases (*n* = 20) and coloured by velocity. **g**, Pearson (*r,* linearity) versus Spearman correlation (𝜌, monotonicity) between negative stimulus distance and HSE membrane potential (blue) or spatially integrated T4a calcium signals (red) for individual approach phases (*n* = 20).

To compare responses across both visual hemifields, we generated a mirrored version of the visual stimulus. This generated population responses for the right visual field and corresponding mirrored estimates for the left. Across both hemifields, the spatial distribution of T4a population activity followed the geometry of the scene (Fig. 4c). Responses were restricted to regions containing visible landmarks and increased as surfaces approached the fly (Extended Data Fig. 6) matching the structure of the environment.

Along the virtual trajectory, neural activity closely followed changes in distance during individual approach episodes (Fig. 4d). During monotonic approaches, decreasing wall distance was accompanied by gradual changes in both HSE membrane potential and spatially integrated T4a calcium signals. This pattern was consistently observed across approach phases for both mirrored scenes from the contralateral and original scenes from the ipsilateral hemifield.

Across monotonic approach phases, neural responses closely tracked wall distance, capturing both the relative decrease in distance over time (Fig. 4e) and the absolute distance, spanning slightly more than ±2 standard deviations of the stimulus range (Fig. 4f). Notably, these responses were observed despite substantial variation in velocity along the trajectory (Fig. 4f and Extended Data Fig. 5e).

Pearson correlation coefficients between neural activity and object distance were high across approach phases (HSE membrane potential: *r* = 0.72 ± 0.24; T4a calcium: *r* = 0.80 ± 0.30; mean ± s.d.). Spearman rank correlations were similarly strong (HSE membrane potential: *ρ* = 0.68 ± 0.30; T4a calcium: *ρ* = 0.78 ± 0.31), indicating a robust monotonic and approximately linear relationship between neural responses and object distance (Fig. 4g).

We next asked whether calcium responses are better explained by distance, velocity, or optic-flow-like signals (Extended Data Fig. 7). Negative distance (−*d*) provided the strongest predictor (R² = 0.74 ± 0.21), whereas velocity also explained substantial variance (R² = 0.67 ± 0.26). Notably, velocity was negatively related to calcium activity (Fig. 4f), with higher velocities associated with lower responses, opposite to the positive velocity dependence expected for optic flow. Inverse distance and optic-flow-like predictors (1/*d*, v/*d*) performed markedly worse (Extended Data Fig. 7).

### Distance-dependent flight control requires motion-based distance estimation

Given the robust and consistent correlations observed previously, the spatial integration of elementary motion detector signals is expected to carry meaningful information about environmental structure. If so, this should manifest in measurable functional consequences: First, population activity combined with the flight trajectory should be sufficient to reconstruct environmental geometry. Second, loss of motion detection should impair distance perception.

To test the first prediction, we transformed T4a calcium signals into distance estimates and mapped them onto allocentric coordinates along the flight trajectory (Fig. 5a and Extended Data Fig. 8). This reconstruction revealed a coherent representation of environmental geometry, with increased point density at the positions of nearby walls. Notably, this structure became more pronounced with spatial integration of the signal. In contrast, a null model in which all directional variation was replaced by a constant mean distance (see Methods) yielded a reconstruction dominated by local sampling density along the trajectory, rather than the geometry of the environment.

**Fig. 5.**
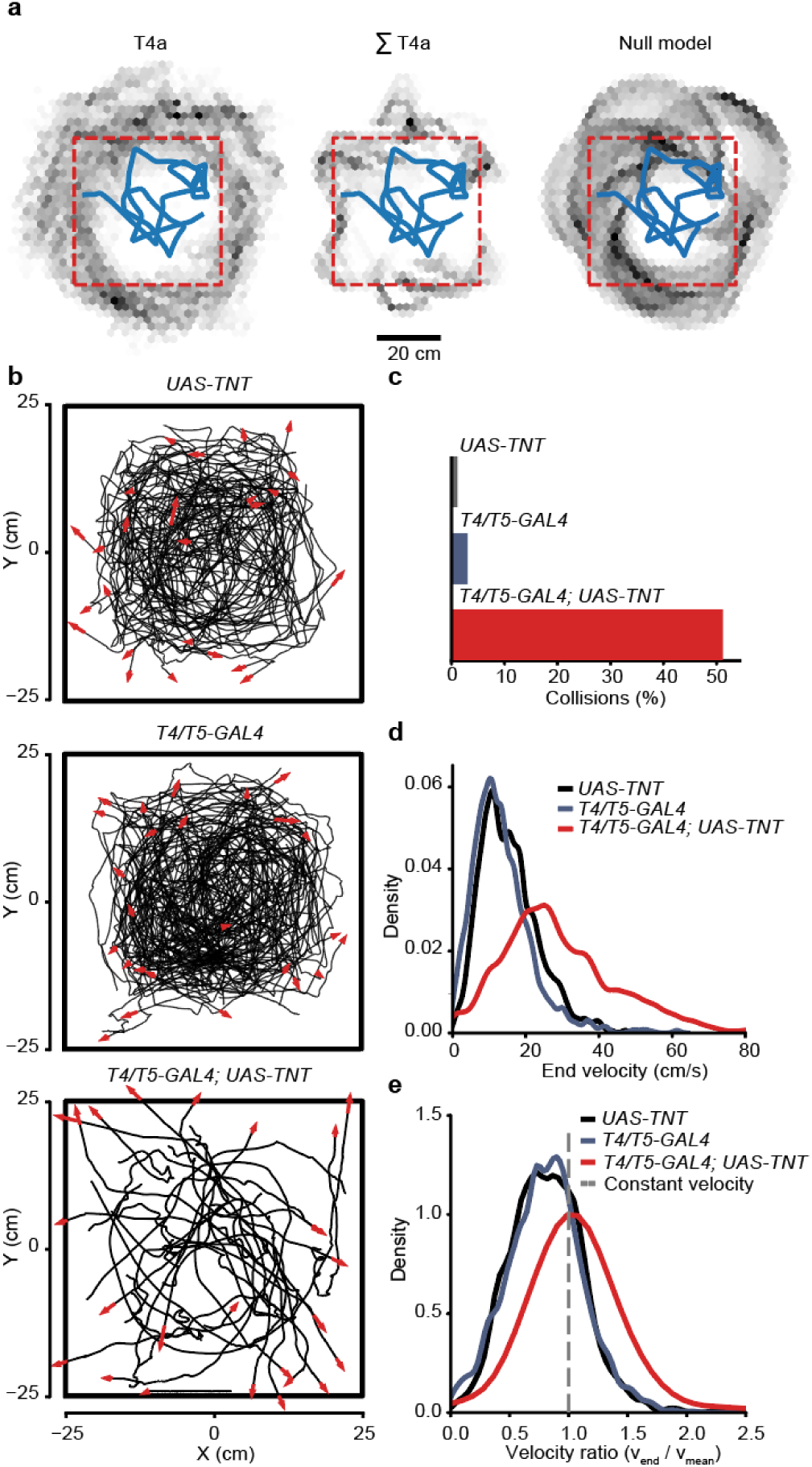
Distance-dependent flight control requires motion vision. **a**, Reconstruction of environmental geometry from population activity. Spatial maps obtained by transforming neural activity-based distance estimates into allocentric coordinates along the flight trajectory. Left, reconstruction based on local T4a activity. Middle, reconstruction after spatial integration across the population (ΣT4a). Right, null model lacking structured distance information. Blue line, flight trajectory; dashed red square, arena boundaries; grey levels indicate point density, jointly normalised across panels. **b**, Projections of the 25 longest free-flight trajectories recorded in a cubic arena with checkerboard-patterned walls of parental control flies (top and centre) and flies expressing tetanus toxin light chain in T4 and T5 neurons (bottom). Red arrows indicate flight direction at trajectory termination; arrow lengths correspond to instantaneous velocity. **c**, Fraction of trajectories terminating in wall collisions for the three genotypes. **d**, Distributions of flight velocities at trajectory termination. Kruskal–Wallis test followed by Dunn’s multiple-comparisons test detected a significant difference in end velocities between control flies and flies with silenced T4/T5 neurons (*P* = 5.42 × 10⁻¹³⁷). **e**, Distribution of the ratio between end velocity (*v*ₑₙd) and mean flight velocity (*v*ₘₑₐₙ). A value of 1 (dashed line) corresponds to constant velocity. Control flies reduced their speed before trajectory termination, whereas flies with silenced T4/T5 neurons exhibited velocity ratios closer to or above unity.

To test the behavioural consequences and separate causation from correlation, we analysed free-flight trajectories of flies in which the outputs of motion-sensitive T4 and T5 neurons were blocked, and compared them with those of parental controls (Fig. 5b and Supplementary video 3). Three-dimensional trajectories were recorded in a cubical enclosure with checkerboard-patterned walls using five cameras^24^. Parental control flies sustained uninterrupted flight for extended distances within the arena, routinely travelling several metres without collisions. Flies that expressed tetanus toxin light chain specifically in T4 and T5 neurons failed to steer clear of walls, resulting in a significant increase in collisions with the arena boundaries (Fig. 5b). More than 50% of flight paths in flies with silenced T4 and T5 neurons terminated in collisions with the wall (Fig. 5c). A Chi-square test of independence revealed a significant relationship between genotype and collision frequency (χ² = 1194.86, *P* = 3.46 × 10⁻²⁶⁰). Control flies reduced their velocity as they approached the end of a flight trajectory, whereas flies lacking functional T4 and T5 neurons failed to decelerate in response to decreasing distance (Fig. 5d, e). Without functional motion detectors, they crashed into obstacles at full speed.

## Discussion

A large body of research describes in detail how individual neurons compute the direction of visual motion^25,26^, but studies that test the behavioural relevance of this computation under naturalistic conditions are scarce.

Traditionally, elementary motion detectors are considered as local direction-selective units that feed into circuits for course stabilization^27,28^. Our results reveal a broader role: Neural activity across a retinotopic array of elementary motion detectors contains information about the geometric structure of the environment, and flies rely on this visual motion-derived depth map to navigate and avoid collisions (Fig. 5).

During translational optic flow, depth information emerges directly from the spatial organization and intrinsic response properties of motion detectors. A shift in coding regimes enables robust proximity signals across scales. At large distances, small retinal displacements are reflected in the graded responses of individual detectors (Fig. 2a, b). As objects approach and local responses saturate, distance information becomes encoded in the spatial distribution of activity across the detector array (Figs. 2 and 3b, c). Spatial integration does not create this signal; it reads out a geometry-dependent recruitment pattern that is already present in the early representation of motion information. Proximity information therefore emerges from the interaction between retinal geometry, motion detector activity, and population structure, without the need to adapt the underlying motion computation.

These findings distinguish motion-derived proximity from time-to-collision computations, which do not require explicit knowledge of distance. Instead, they rely on the temporal derivative of retinal image expansion and image size on the retina to predict the time of impact^29–31^. This is often invoked as the primary motion-based variable guiding avoidance or landing. However, neurons that compute the time to collision typically respond to small looming stimuli^32^, and are inhibited by wide-field motion such as flow fields generated by egomotion^33^. Our data indicate that early motion-processing circuits provide a geometry-derived proximity signal that scales monotonically with distance across a broad behavioural range and remains locally stable despite fluctuations in translational speed (Figs. 3d and 4f). This population-based proximity signal persists under naturalistic visual motion statistics and is behaviourally relevant. During replay of natural flight trajectories, spatially integrated T4a activity and responses of postsynaptic HSE cells increase systematically as animals approach surfaces (Fig. 4). In freely flying flies, intact motion vision supports deceleration as they approach obstacles. Silencing T4 and T5 neurons abolishes this distance-dependent speed regulation and leads to frequent collisions with obstacles (Fig. 5b–e). Early motion pathways are therefore not merely correlated with environmental proximity; they are required to translate retinal motion into adaptive control of flight speed.

More broadly, our findings suggest that spatial structure can be derived from retinal dynamics without invoking dedicated depth-specific circuitry. Simple spatial integration within early sensory pathways may provide a general strategy by which nervous systems extract geometric information from self-generated motion. Environmental layout is embedded in the distributed activity patterns of primary motion detectors.

## Methods

### Fly husbandry

*Drosophila melanogaster* were cultivated on a cornmeal, molasses and yeast medium under a 12 h–12 h light–dark cycle at 25 °C and 60% relative humidity.

For generating the *T4a/T5a-GAL4* driver line, the chosen enhancer sequence (chr2L: 8,782,850-8,783,701, GCF_000001215.4) corresponds to a shortened version of the *GMR39H12* enhancer^1^, retaining only the region of high chromatin accessibility in T4 and T5 neurons as assessed by ATACseq^2^. The enhancer sequence was synthesized by Genewiz (Azenta) and inserted into the *pBPGUw* plasmid backbone^3^ (RRID: Addgene_17575) after restriction digest with ZraI and NaeI. Embryo injections for PhiC31 integrase-mediated transgenesis at the attP40 landing site were performed by BestGene, Inc. The expression pattern of the newly generated driver line was assessed in flies additionally bearing a GFP construct (RRID: BDSC_32188) of the genotype *w+/w–; P{T4a/T5a-GAL4}attP40/P{10XUAS-IVS-mCD8::GFP}su(Hw)attP5*. In contrast to the original driver line *P{GMR39H12-GAL4}attP2* that targets all T4/T5 subtypes^1,4^, this driver line only targets the a-subtypes (Extended Data Fig. 1a, b).

For calcium imaging experiments, flies from the *T4a/T5a-GAL4* driver line were crossed to *GCaMP8m*-bearing flies^5^ (RRID: BDSC_92590) to obtain flies with the genotype *w+/w–*; *P{T4a/T5a-GAL4}attP40*/+; *P{20XUAS-IVS-jGCaMP8m}VK0005*/+.

For HSE electrophysiology experiments, flies with the genotype *w+/w–; P{10XUAS-IVS-mCD8::GFP}su(Hw)attP5; P{GMR81G07-GAL4}attP2* (RRID: BDSC_32188 and RRID: BDSC_40122) were used^1^.

For characterizing naturalistic flight patterns in a closed environment, we used wild-caught flies (‘Luminy’). To generate motion-blind flies, we used a split-GAL4 driver line (combination of RRID: BDSC_70685 and BDSC_72763) to drive the expression of tetanus toxin light chain^6^ (RRID: BDSC_28837) in all T4 and T5 cells in the optic lobe, resulting in the following genetic line: *w+/w–; P{R42F06-p65.AD}attP40/P{UAS-TeTxLC.tnt}E2; P{VT043070-GAL4.DBD}attP2/+.* Both parental controls were obtained by crossing the respective lines to CantonS flies, thus obtaining *w+/w–; P{R42F06-p65.AD}attP40/+; P{VT043070-GAL4.DBD}attP2/+* for the split-GAL4 control and *w+; P{UAS-TeTxLC.tnt}E2/+* for the UAS control.

### Histology

Fly brains (1–6 days after eclosion) were dissected in phosphate-buffered saline (PBS; 137 mM NaCl, 3 mM KCl, 8 mM Na2HPO4, 1.5 mM KH2PO4, pH 7.3), fixed for 30 minutes in 4% (w/v) paraformaldehyde in PBS with 0.1% (v/v) Triton X-100 and washed in PBS with 0.3% (v/v) Triton X-100 (PBT). The brains were blocked overnight in PBT with 10% normal goat serum (NGS) and incubated with primary antibodies mouse anti-nc82 (1:20, DSHB, AB2314866) and chicken anti-GFP (1:500, Rockland, 600901215). After three one-hour washes in PBT at room temperature, the brains were incubated with secondary antibodies goat anti-mouse Alexa Fluor 568 (1:500, Invitrogen, A-11004) and goat anti-chicken Alexa Fluor 488 (1:500, Invitrogen, A-11039). All antibodies were diluted in PBT with 5% (v/v) NGS. Both incubations lasted at least 48 hours at 4 °C. After antibody incubations, the brains were washed in PBT at 4 °C overnight and mounted in SlowFade Gold Antifade Mountant (Invitrogen, S36937). Micrographs were acquired on a Leica Stellaris 5 laser scanning confocal microscope using an HC Plan-apochromat glycerol-immersion objective (63x/1.3 NA) and Leica Application Suite X software (all Leica), and processed using the Fiji distribution of ImageJ^7^.

### Visual stimulation and scene generation

#### Panoramic projection display

Visual stimuli were presented using a projection-based panoramic screen in which a digital projector illuminated the inner surface of a hemispherical bowl surrounding the fly. This geometry allowed visual textures of custom geometry to be displayed across the visual field, embedding the animal in a virtual visual environment, as previously described^8^. Visual scenes were defined in spherical coordinates with a fixed geometric correspondence between projector pixels and visual angles at the fly’s position (0.5° per pixel).

A ViewSonic M2e projector was used for all experiments, providing 1,280 × 720-pixel resolution at a refresh rate of 120 Hz. Luminance uniformity and contrast were comparable to those reported previously^8^. The stimulation hardware was redesigned using a combination of 3D-printed components and standard optomechanical elements (Thorlabs), reducing system complexity while preserving geometric precision.

To enable simultaneous two-photon calcium imaging, the visual stimulation system was modified to minimize interference with fluorescence detection. The blue and red LED channels of the projector were disabled, and the projected light was spectrally filtered using long-pass filters (Thorlabs FEL0550 and FGL550). The stimulation device was additionally enclosed in a light-shielding housing to prevent stray light contamination.

#### Adaptations for in vivo electrophysiology

For in vivo whole-cell patch-clamp recordings, the visual stimulation system was configured to minimize electromagnetic interference with electrophysiological recordings. The stimulation apparatus was shielded using conductive metal foil, and all electrical components were grounded to reduce line noise and pickup issues. No optical filtering or light-shielding enclosure was used during electrophysiological recordings.

#### Visual scene renderings

Visual stimulus videos were generated using Blender (v4.2). Scenes were modelled in three dimensions and adapted to match the experimental geometry. A virtual camera representing the observer was placed within the scene and configured to render panoramic videos covering 360° × 180° of visual space at a resolution of 720 × 360 pixels, corresponding to an angular resolution of 0.5° per pixel. Rendering was performed using an equidistant cylindrical projection. No contrast-enhancing or other image-altering postprocessing was applied.

Motion stimuli were generated by animating the virtual observer to traverse the scene at constant speed and rendered using the cycles rendering engine in Blender. Stimuli based on recorded flight trajectories were rendered at 100 Hz, matching the temporal resolution of the original recordings. Constant velocity stimuli were rendered at 60 Hz. To prevent unwanted flicker responses, rendering noise was reduced using OpenImageDenoise (Intel^®^) in Blender, guided by auxiliary render passes.

For free-flight replay and the digital twin of the free-flight arena, only patterned surfaces were rendered. The floor and ceiling were represented as uniform mid-grey surfaces, and transparent acrylic elements originally used to construct the arena were omitted to reduce scene complexity. Flight trajectories were imported using custom Python v.3.7 (Python Software Foundation) scripts. Measured positional and kinematic parameters, including position and velocity, were used to animate the virtual camera.

Viewing direction was reconstructed from the trajectories using a simplified active gaze model, rather than assuming passive motion. No smoothing, projection, or additional adaptation of the trajectory data was applied. Head and eye movements were not measured and were therefore not incorporated into the stimulus reconstruction. In the active gaze model, the virtual observer was allowed to rotate only during saccadic flight segments. During inter-saccadic intervals, gaze direction was assumed to remain fixed and aligned with the instantaneous flight direction. This assumption reflects the well-characterized flight behaviour of flies^9–12^, in which gaze is actively stabilized between saccades through mechanosensory and visual feedback, minimizing rotational and emphasizing translational optic flow.

#### Synchronization

Visual stimulus presentation was synchronized with neural recordings using a photodiode-based trigger system (Texas Instruments, OPT101) in both two-photon calcium imaging and electrophysiological experiments. The projector generated light flashes at stimulus onset, during playback, and at stimulus termination. These signals were detected by a photodiode positioned outside the projection screen, such that trigger detection did not interfere with the visual stimulation presented to the animal.

The photodiode output was recorded as an Independent analogue input channel and acquired using the same data acquisition pipeline as the corresponding neural signals (calcium imaging frames or electrophysiological recordings). This provided a common and consistent temporal reference for precise alignment of visual stimulus timing with neural data across both experimental modalities.

### In vivo two-photon calcium imaging

#### Two-photon microscopy

Functional calcium imaging of T4a neurons in the right optic lobe was performed in female flies using microsurgical procedures adapted from Maisak et al.^13^ Flies were cold-anesthetized and mounted on an acrylic glass holder with beeswax to immobilize thorax and legs. The head was gently bent to expose the posterior surface and inserted into a custom-cut opening in aluminium foil clamped into a recording chamber filled with external saline (pH 7.3) containing 5 mM TES, 103 mM NaCl, 3 mM KCl, 26 mM NaHCO3, 1 mM NaH2PO4, 1.5 mM CaCl2, 4 mM MgCl2, 10 mM trehalose, 10 mM glucose and 7 mM sucrose (280 mOsM, equilibrated with 5% CO2 and 95% O2). A small window was cut into the right side of the head capsule, and muscles, adipose tissue, and trachea were removed to expose the optic lobe.

Imaging was performed on a custom-built two-photon laser-scanning microscope^14^ equipped with a 40× water-immersion objective (0.80 NA; IR-Achroplan, Zeiss). Fluorescence excitation was provided by a mode-locked Ti:sapphire laser (Mai Tai HPDS; Spectra-Physics) tuned to 910 nm. Excitation power at the sample was adjusted to 10–20 mW using a power attenuator (CCVA-PR-CD; Spectra-Physics). Emitted fluorescence was detected with a photomultiplier tube (H10770PB-40; Hamamatsu) positioned behind a 509/22 nm band-pass filter (BrightLine).

Two-photon images were acquired using ScanImage 3.8 (Vidrio Technologies) running in MATLAB R2013b (The MathWorks). Two-photon calcium imaging was performed in T4a neurons expressing GCaMP8m. Calcium signals from T4a dendrites were recorded at a spatial resolution of 32 × 32 pixels and a frame rate of 25.2 Hz.

#### Visual stimulation protocol and experimental timeline for two-photon calcium imaging

All experiments followed a standardized visual stimulation protocol (Extended Data Fig. 4). Individual protocols differed in the duration and content of the video stimuli, but shared a common structure and were constrained to a total duration of less than 10 min per recording to minimize slow tissue drifts during two-photon imaging.

Each recording began with a binary white-noise stimulus (5 min), which was used exclusively for receptive-field estimation, followed by structured and naturalistic video stimuli. Individual video durations varied across protocols but ranged from 4 to 18 s. Each video stimulus was preceded by a 2-s static pre-stimulus phase, during which the first frame of the video was presented without motion, followed by the motion sequence and a 2-s static post-stimulus phase displaying the last frame of the video before the onset of the next stimulus.

Each video stimulus was presented multiple times within a recording, with a minimum of three repetitions per stimulus. No averaging across stimulus repetitions was performed at this stage.

#### Image registration

Two-photon image sequences were corrected for drift using rigid, translation-only image registration. Following registration, recordings were visually inspected prior to analysis to ensure signal stability and consistent responses over time.

Data quality was further validated using the white noise stimulus. Recordings that did not exhibit reliable correlations with the noise stimulus were automatically excluded from further analysis. All assessments were performed blind to stimulus identity.

#### White-noise stimulation and receptive-field mapping

For receptive field mapping, a binary white-noise stimulus was used, consisting of spatially independent luminance values updated at 25 Hz. The stimulus was defined in visual space and discretized into a grid of 28 × 36 stimulus pixels, corresponding to approximately 5° per pixel in visual angle, with luminance values drawn from a binary distribution (0 or 128).

During white-noise presentation, fluorescence signals were recorded simultaneously across the imaging field. Fluorescence time series were extracted post hoc on a per-pixel basis in imaging space (32 × 32 pixels per imaging plane), without predefined regions of interest. This pixel-wise analysis accommodated the densely interwoven dendritic morphology of T4a neurons and the curvature of the medulla.

Receptive fields were estimated using a tensor-based reverse-correlation approach implemented in the frequency domain. Stimulus sequences and fluorescence signals were represented as multidimensional tensors, enabling parallel computation of spatiotemporal stimulus–response kernels across spatial locations. Fluorescence signals were temporally resampled to match the stimulus time base, and low-frequency trends were removed prior to analysis.

Correlations were computed using fast Fourier transforms (FFTs), followed by inverse FFTs to recover spatiotemporal receptive-field kernels in the time domain. This tensorized formulation allowed FFT operations to be applied simultaneously across spatial dimensions. The analysis was implemented in JAX (Google), enabling automatic vectorization and parallelization. This implementation reduces computational complexity from *O* (*T*^2^) to *O* (*T log T*). In practice, FFT-based correlation yielded a ∼5–6× speedup over time-domain methods, which increased to ∼100× when combined with tensorization and JAX-based parallelization, reducing computation times from hours to seconds.

Receptive field locations were estimated by identifying the visual stimulus positions that reliably evoked responses in each imaging pixel. Spatial receptive fields were computed for each imaging pixel by averaging spatiotemporal kernels over a time window of 0.08 to 0.24 s following stimulus updates. Resulting maps were spatially smoothed and *z*-scored, and pixels were classified as responsive if the peak response exceeded a conservative threshold of |z| > 5. For responsive pixels, receptive field centres were estimated by fitting a two-dimensional Gaussian to the spatial receptive field map. This fitting was used solely to estimate the preferred location in visual space and not to characterize receptive-field shape.

Pixels with identical receptive field locations were grouped automatically into ROIs in imaging space. ROI definitions were derived exclusively from the white-noise mapping data and were independent of responses to structured video stimuli. Receptive field kernels were averaged within each ROI for subsequent analyses.

For each ROI, fluorescence time series were obtained by averaging signals across all included pixels. The centroid position of each ROI in imaging space and its corresponding receptive field location in visual space were determined, and stored together with the fluorescence trace for each imaging plane. These ROI definitions were kept fixed and used consistently across all analyses of structured and naturalistic video stimuli.

#### Stimulus-aligned preprocessing and signal normalization

Imaging data were segmented into stimulus-aligned epochs based on the recorded photodiode trigger signals. Only frames corresponding to active stimulus presentation were included in further analyses. Baseline fluorescence (*F*₀) was computed separately for each ROI and stimulus presentation as the mean fluorescence during the pre-stimulus baseline period defined relative to stimulus onset. Relative fluorescence changes were calculated as *ΔF/F* = (*F*− *F*₀)/*F*₀. No temporal or spatial averaging across stimulus repetitions was performed during preprocessing. For each stimulus presentation, a separate *ΔF/F* trace was stored together with its corresponding stimulus metadata.

#### Multi-position and multi-fly data acquisition

For each fly, receptive field mapping and ROI definition were performed independently for every imaging stack. Pixel-wise receptive fields were estimated from the white-noise stimulus, and imaging pixels with identical receptive field locations were grouped into regions of interest (ROIs). Only ROIs that met the predefined quality criteria were retained. Each ROI was assigned a preferred position in stimulus space based on its receptive field centre and constituted the fundamental unit for all subsequent analyses.

To increase spatial coverage of the medulla within individual flies, multiple imaging stacks were recorded sequentially from different positions (in X, Y and Z) in the same animal. All stacks within a fly were recorded using the same visual stimulation protocol, but at different time points during the experiment.

Across flies, data were pooled exclusively based on receptive field location in stimulus space. ROIs from different animals with matching receptive field positions were assigned to the same stimulus coordinates, such that population responses at a given location represent pooled measurements across ROIs, imaging positions and flies. All flies contained in a given dataset were presented with the identical visual stimulation protocol. Comparisons were therefore restricted to within-dataset analyses; responses obtained under different stimulation protocols were not compared directly.

Dataset 1 (Figs. 3e and 4) comprised 80 imaging stacks obtained from 9 flies. Across all stacks, a total of 1065 valid ROIs were included. On average, imaging stacks contained 13.3 ± 5.1 ROIs (mean ± s.d.). The dataset comprised 227 locations, corresponding to an angular coverage of 5,675 deg², equivalent to a circular visual field with a diameter of ∼85°.

Dataset 2 (Figs. 1, 2, and 3d, f) comprised 107 imaging stacks obtained from 12 flies. Across all stacks, a total of 1124 valid ROIs were included. On average, imaging stacks contained approximately 10.5 ± 5.5 ROIs (mean ± s.d.). The dataset comprised 223 locations, corresponding to an angular coverage of approximately 5,575 deg², equivalent to a circular visual field with a diameter of ∼84°.

#### Stimulus–response maps

Within each dataset, population activity was expressed in stimulus coordinates by combining responses across imaging positions and flies using the receptive field mapping described above. ROI fluorescence signals were assigned to their preferred stimulus locations and pooled across recordings to generate spatiotemporal stimulus–response maps. These maps describe population activity as a function of visual position and time and enable direct comparison with stimulus-derived features defined in the same coordinate space.

At each stimulus location, signals from multiple ROIs, imaging positions, and flies were averaged. Stimulus–response maps provided a representation of neural population activity aligned to visual coordinates and formed the basis for all subsequent analyses relating neural responses to local stimulus properties such as luminance, distance, and motion dynamics.

#### Visualization and data export

Stimulus–response maps were used to visualize population activity in stimulus coordinates (Fig. 1e, 2a and 4b, Supplementary video 1 and 2). Time-resolved spatial maps, XT representations (visual position × time), and overlay visualizations combining neural activity with the corresponding stimulus frames were derived directly from these maps. For the XT representation in Fig. 1e and 4c, activity was averaged across defined bands in visual space orthogonal to the axis of motion.

All visualizations were generated using identical processing and scaling parameters across datasets. Images, XT plots, and stimulus-aligned response videos were stored together with the underlying numerical data to enable reproducible analysis and figure generation. These outputs served as the basis for quantitative analysis and figure preparation.

### In vivo electrophysiology

#### Patch-clamp

Whole-cell patch-clamp recordings were carried out in vivo with female flies aged 2–12 h post-eclosion. They were cold-anaesthetized and fixed to a laser-cut polyoxymethylene mount with soft thermoplastic wax (Agar Scientific). The fly head was partially submerged in extracellular solution (pH 7.3; 280 mOsM, equilibrated with 5% CO₂ and 95% O₂) containing 103 mM NaCl, 3 mM KCl, 26 mM NaHCO3, 1 mM NaH2PO4, 1.5 mM CaCl2, 4 mM MgCl2, 10 mM trehalose, 10 mM glucose, 5 mM TES and 7 mM sucrose. The right dorsal optic lobe and the central brain were exposed by surgically removing cuticle, trachea, and adipose tissue. Patch pipettes (5–7 MΩ) with outer and inner diameters of 1.5 mm and 1.17 mm, respectively, were made from borosilicate glass capillaries using a P-97 (Sutter Instruments) micropipette puller. Pipettes were filled with intracellular solution containing 10 mM HEPES, 140 mM potassium aspartate, 1 mM KCl, 4 mM MgATP, 0.5 mM Na3GTP, 1 mM EGTA (265 mOsM, pH 7.3). A combination of bright-field and epifluorescence microscopy was used to visually find green fluorescent somata using an Axio Scope.A1 microscope (Zeiss) equipped with a ×60/1.0 NA water-immersion objective (LUMPLFLN60XW, Olympus) and an LQ-HXP 120 light source (Leistungselektronik Jena). Transillumination was achieved by attaching a white LED (MCWHD5, Thorlabs) to a light guide, the far end of which was positioned beneath the holder in the left hemisphere of the fly. The perineural sheath was cut using a micropipette filled with extracellular solution (see above) to expose somata. The *GMR81G07-GAL4* driver line labelled all three horizontal-system (HS) neuron subtypes. Individual HSE neurons were identified anatomically based on the position of their somata within the optic lobe.

Signals were recorded using a MultiClamp 700B amplifier, low-pass filtered and digitized at 10 kHz with a Digidata 1550B and controlled via pCLAMP 11 software (all from Molecular Devices). Data were corrected for the liquid junction potential (12 mV). Analysis was performed with software in Python v.3.9 (Python Software Foundation) using NumPy v.2.3, Pandas v.2.3, SciPy v.1.16, Matplotlib v.3.10 and pyABF v.2.3 (https://pypi.org/project/pyabf/). Resting potential was defined as the mean membrane voltage during the non-stimulus period of the first sweep. Only cells with resting potentials more negative than −25 mV and voltage responses to the translation-edge stimulus exceeding 3 mV were included.

#### Visual stimulation protocol and experimental timeline for in vivo electrophysiology

Patch-clamp recordings followed the same standardized visual stimulation protocol used for two-photon imaging. Stimulus order during electrophysiological recordings was not randomized, and stimuli were presented in a fixed sequence. Stimuli were repeated for as long as stable whole-cell recordings could be maintained. Recordings were included based on the stability of the baseline membrane potential, which was required to remain more negative than −25 mV throughout the recording. Furthermore, cells were only included if their response to the translational edge stimulus exceeded 3 mV in amplitude.

#### Stimulus pre-processing

Electrophysiological recordings were analysed offline using custom routines written in Python v.3.9 (Python Software Foundation). Membrane potential traces were extracted from ABF files, and photodiode trigger signals were used to align recordings to stimulus onset and offset. For each stimulus presentation, voltage traces were segmented into stimulus-aligned epochs including a 1-s pre-stimulus baseline period. Baseline voltage was subtracted using the mean membrane potential during this interval. Stimulus identity and parameters were obtained from synchronized metadata files and used to group trials belonging to the same stimulus condition. Within each condition, single-trial responses were averaged to obtain mean membrane potential traces for HSE neurons. For comparisons across recordings, stimulus-aligned responses were pooled based on identical stimulus metadata and summarized as mean responses across trials. In contrast to T4a calcium imaging, receptive fields were not mapped for HSE recordings. Consequently, potential shifts in receptive-field location arising from the experimental preparation cannot be excluded.

### Free flight experiments

Details of the free-flight arena and trajectory reconstruction were described previously^15^. Key experimental procedures are summarized below.

#### Flight arena

A transparent enclosure with the dimensions 50 × 50 × 30 cm made of acrylic plastic (Evonik, Plexiglas XT) was used. To facilitate optical tracking, a custom LED-array of high-intensity infrared lights (830 nm, Roithner Laser Technik, GmbH) along with a diffuser were placed under the arena. Visual stimulation was administered via 4 monitors (144 Hz, ASUS, VG 248QE) surrounding the vertical walls of the arena. The monitors were controlled via NVIDIA 3D Vision Surround Technology on Windows 7 (64-bit). The visual stimulus consisted of a static checkerboard pattern (5 × 5 cm) alternating light (5 cd m^−2^) and dark squares (2 cd m^−2^). Stimuli were displayed using Panda3D and Python 2.7.

#### Fly preparation

Female flies that were younger than 48 hours were selected under brief CO2 anaesthesia. They were left to recover for 24 hours in a new vial with standard food, after which they were transferred to an empty vial where they were starved for 4 hours prior to the start of the experiment. The experiments were started at the same time of the day and were performed overnight. The ambient temperature inside the arena was 28 °C.

#### Flight path tracking

Flies were recorded using five cameras (FLIR Inc., CM3-U3-13Y3M-CS) equipped with near-infrared long-pass filters (Thorlabs Inc., FGL 780M) and machine vision lenses (focal length 6 mm, Thorlabs Inc., MVL6WA). The cameras were all connected to one computer, and synchronised by a TTL pulse generator operating at 100 Hz. Each pulse resulted in the acquisition of 5 individual 640 × 512 pixel single-channel matrices. Calibration was performed periodically using the single-step method described by Li et al.^16^ Each calibration consisted of 100–200 synchronised images of a random pattern. We used the standard pinhole camera model with radial distortion provided in the MATLAB toolbox (https://github.com/prclibo/calibration-toolbox).

For each frame, moving objects were detected independently in each camera view using background subtraction with exponential updating, followed by noise suppression and blob-centre extraction. The resulting two-dimensional coordinates were triangulated across the five camera views to obtain the three-dimensional position of each fly at every time step.

We performed a per-frame triangulation of all five concurrent two-dimensional coordinates to determine the position of the flies in the 3D space of the arena at each time step. For multiple flies changing position at the same time, we achieved this using the standard singular value decomposition^17^. Individual flight trajectories were reconstructed using a linear Kalman-filter framework with three-dimensional position as the observation variable. Frame-to-frame assignment of observations to existing trajectory estimates was solved using an optimal assignment procedure^15^. If no new observations were added for 20 new frames, the instance was terminated. The resulting trajectory was then saved. Detection and trajectory reconstruction were performed offline in Python 2.7 using the FilterPy library (https://filterpy.readthedocs.io/en/latest/).

#### Trajectory selection for free-flight replay

From the wild-caught flies’ dataset, the 30 longest flight trajectories were selected based on total path length. Candidates were evaluated using four criteria: Z-position variance (vertical stability; ranked ascending), deviation of mean Z from 15 cm (target height consistency; |mean Z – 15 cm|, ascending), average length of straight segments (descending), and deviation of average speed from the population median (locomotor representativeness; |speed − median|, ascending). Low vertical variability ensured stable stimulus placement within the arena and constrained motion to the horizontal axis relevant for T4a direction-selective neurons, while target height consistency maintained central stimulus alignment. Trajectories were ranked per metric, and ranks were summed to obtain a composite score. Among the highest-ranked candidates, trajectory “target_1056_path_straight.csv” was selected because it combined low vertical variability with pronounced depth modulation and broad depth sampling, together with a wider speed regime resulting in increased spatial coverage.

#### Free-flight dataset and collision analysis

Free-flight experiments were conducted across three genotypes: *UAS-TNT*, *T4/T5-GAL4*, and *T4/T5-GAL4*; *UAS-TNT*. A total of 73 flies were recorded, yielding 3,214 flight trajectories (*T4/T5-GAL4*; *UAS-TNT*: 640 trajectories from 35 flies; *UAS-TNT*: 1,263 trajectories from 18 flies; *T4/T5-GAL4*: 1,311 trajectories from 20 flies).

Trajectories were classified as collisions if flies either directly reached the arena walls or, based on their instantaneous velocity at the end of the trajectory, were projected to hit the wall within the next 100 ms. Using this criterion, the number of collision events was 331/640 trajectories for *T4/T5-GAL4; UAS-TNT*, 14/1,263 for *UAS-TNT*, and 41/1,311 for *T4/T5-GAL4*.

### Algorithmic motion detector model

We implemented a fully specified, feedforward three-arm motion detector operating on spatiotemporal luminance sequences. Visual input (360 × 720 pixels, 60 Hz, 0.5°/pixel) was sampled along the horizontal axis using Gaussian spatial filters (σ = 6.2°, 9°, and 6.2° for leading, central, and trailing arms, respectively) spaced at 5° intervals. This produced local intensity time series that were normalised to the range [−1,1].

The leading and trailing arms each applied a first-order low-pass filter with a time constant *τ* of 0.25 s. The central arm formed a band-pass filter with a low-pass (*τ* = 0.15 s) and a high-pass (*τ* = 1.5 s) component. Simulations were performed with a time step of 16.6 ms. Temporal filtering was followed by half-wave rectification with zero threshold. Motion signals were computed from neighbouring spatial columns by combining arms asymmetrically: the rectified difference between leading and trailing arm activity (plus a constant bias of 0.2) was multiplicatively gated by the central arm. For simplicity, the model was implemented in one dimension and applied only along a single horizontal slice of the visual field (elevation = 0°).

### Trajectory-based reconstruction model

To relate motion-derived T4a signals to spatial structure, we implemented a trajectory-based reconstruction model. Angular distance profiles were computed over discrete 5° bins and processed using different spatial integration schemes: no smoothing, Gaussian smoothing (∼60°), and hemispheric integration, in which left and right visual fields were averaged independently.

T4a responses were normalised per recording and mapped to physical distances using a linear scaling with an upper bound of 42 cm, corresponding to the 90^th^ percentile of distances observed in the real dataset, providing a robust estimate of maximal range (*d* = 42 – T4a × 42). For each time point, angular distance estimates were projected into world coordinates using the animal’s position and heading, yielding point clouds of inferred object locations.

As a null model, we used an identical reconstruction pipeline but replaced the spatially structured distance signal with its global mean, resulting in a constant radial estimate without directional information. This produced circular reconstructions that served as a baseline for comparison with structure-preserving models.

## Data availability

The datasets generated and analysed during the current study are available from the corresponding author upon reasonable request. Source data underlying the figures are provided with this paper will be made available upon publication at DOI: 10.5281/zenodo.20049664.

## Code availability

Custom code used for stimulus generation, data processing, and analysis is available from the corresponding author upon reasonable request. Example applications, including routines for ROI extraction and supplementary material for figure generation, will be made available upon publication at https://github.com/stefanprech/Motion-based-depth-estimation-in-Drosophila. Large language models (e.g., ChatGPT-4 and ChatGPT-5, OpenAI) were used to assist with code development. All generated code was reviewed, tested, and validated by the authors.

## Statistics and reproducibility

No statistical methods were used to predetermine sample sizes. Sample sizes are comparable to those generally employed in the field. All experiments were replicated across multiple animals, and data were pooled based on matched stimulus conditions and receptive field locations.

Data are presented as mean ± s.e.m. unless otherwise indicated. Statistical analyses were performed using Python (NumPy, SciPy). Pearson’s correlation coefficient was used to assess linear relationships, and Spearman’s rank correlation coefficient was used to assess monotonic relationships.

For free-flight replay experiments, correlations between neural activity and distance were computed across time points within individual approach segments. Correlation coefficients were calculated separately for each segment and summarized across segments.

For stimulus-driven experiments, correlations between neural responses and stimulus signals were computed on grouped data subsets. Signals were partitioned into independent subsets, and correlation coefficients were calculated for each subset. Reported values correspond to the mean across subsets.

A chi-square test of independence was used to assess differences in collision frequency across genotypes. For comparisons of velocity distributions, a Kruskal–Wallis test followed by Dunn’s multiple-comparisons test was applied. Equivalence testing was performed using two one-sided tests (TOST) with predefined equivalence bounds.

Unless otherwise stated, statistical tests were two-sided. Exact sample sizes (n) and definitions of n (e.g., number of flies, trajectories, segments, or grouped subsets) are provided in the corresponding figure legends. No data were excluded unless predefined quality criteria were not met, including unstable baseline signals or insufficient responses to control stimuli. These criteria were applied consistently across all datasets.

The experiments were not randomized, and investigators were not blinded to allocation during data collection and analysis. However, all analyses were performed using automated pipelines applied uniformly across conditions.

## Supporting information

Supplementary Video 1 Visual stimulus and neural response during constant translation

Supplementary Video 2 Virtual flight replay

Supplementary Video 3 Free-flight experiments

## Acknowledgements

We thank G. Rubin for the *pBPGUw* plasmid, B. Prud’homme, F. Schnorrer, and N. Luis for catching and providing the ‘Luminy’ wild-caught strain, T. Schilling for the development and characterization of the *split-GAL4* line used, and A. Mauss and A. Leonhardt for the development of the free-flight tracking system. We further thank Michael Deistler for discussions. Flies obtained from the Bloomington Drosophila Stock Center (NIH P40OD018537) were used in this study. This work was supported by the Max Planck Society and by the European Research Council Starting Grant 101116996 (TemProDroMe) to L.N.G. Views and opinions expressed are those of the authors only and do not necessarily reflect those of the European Union or the European Research Council Executive Agency.

## Author contributions

S.P. conceived the study, designed experiments, established the virtual reality systems, created the stimuli, built the digital twin of the free-flight arena, analysed all data, and ran model simulations. S.P. and J.H. developed the white-noise-based automated ROI selection algorithm. S.S.-H. performed electrophysiological recordings. B.Z. generated fly lines and performed structural imaging experiments. M.-B.L. acquired behavioural data. J.H. performed calcium imaging experiments. L.N.G. and A.B. supervised the study. The manuscript was written by S.P. with the help of L.N.G. and edited by all authors.

---

**Extended Data Fig. 1.**
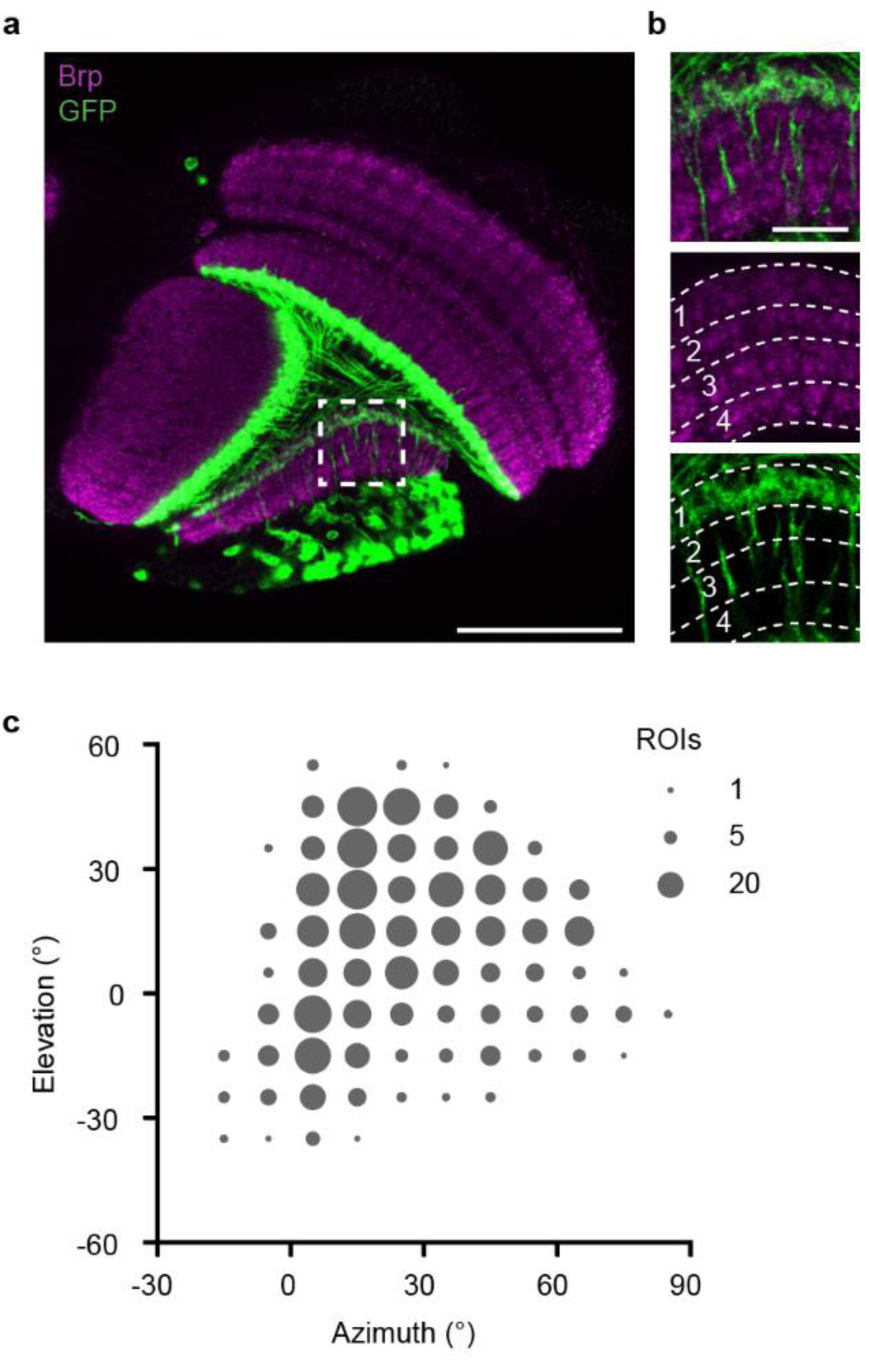
Specificity of the *T4a/T5a-GAL4* line. **a**, Confocal cross-section of the optic lobe of an adult fly expressing GFP (green) under control of *T4a/T5a-GAL4*, counterstained with anti-Bruchpilot (Brp, magenta). The dashed box indicates the lobula plate region shown at higher magnification in **b**. Scale bar, 50 µm. Image representative of 5 biological replicates. **b**, Magnified view of the lobula plate (dashed box in **a**). Top, merged GFP and Brp channels; centre, Brp channel only; bottom, GFP channel only. Dashed lines indicate boundaries between lobula plate layers; numbers denote individual layers. Scale bar, 20 µm. **c**, Density of regions of interest (ROIs) across the reconstructed visual field. Data are binned into 10° × 10° azimuth–elevation bins; circle size indicates the number of ROIs per bin.

**Extended Data Fig. 2.**
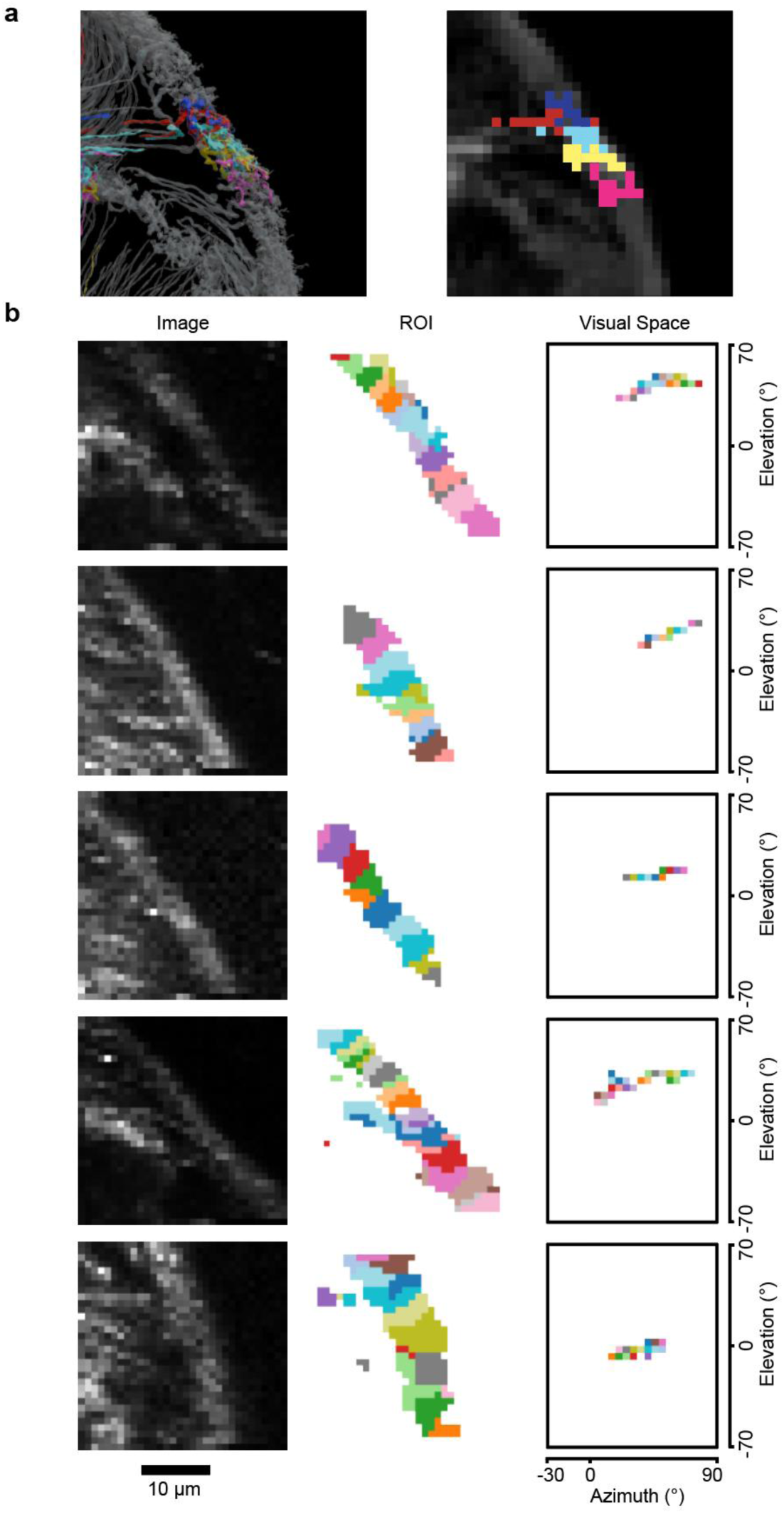
Inferring space from arrays of visual motion-sensitive neurons. **a**, Left, visualization of medulla neuropil with example T4a dendrites highlighted in colour, based on an electron microscopic reconstruction. The view spans ∼30 µm in width and ∼5 µm in depth, comparable to the axial sampling of a two-photon imaging plane. The rendering illustrates the dense and highly intermingled dendritic organization. Right, ROIs manually drawn by a human expert using ground-truth anatomical information. Even with access to detailed structural data, defining accurate and non-overlapping ROIs is challenging due to the strong dendritic overlap. **b**, Examples of ROI selection in different regions of the medulla. Left, example two-photon image patches. Based on these images alone, it is not possible to reliably define ROIs without prior knowledge of the underlying neuronal structure. Centre, ROIs automatically identified using receptive field mapping following white-noise visual stimulation. Right, corresponding receptive field locations projected into visual space in the virtual reality arena (azimuth and elevation).

**Extended Data Fig. 3.**
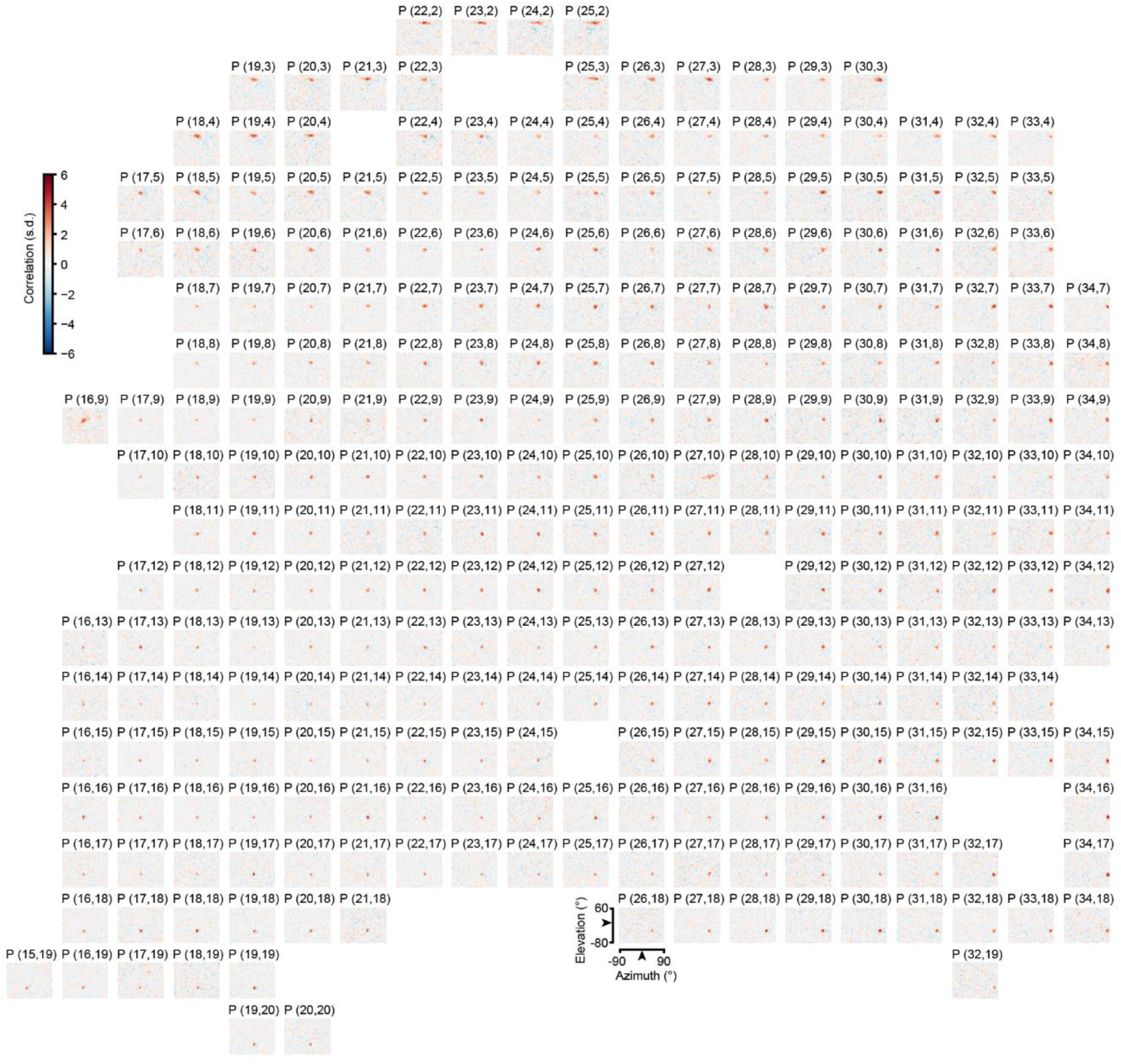
Spatial receptive fields of T4a neurons. Grid-based visualization of receptive field kernels reconstructed for individual ROIs across the visual field based on white-noise stimulation. Each panel P (x, y) corresponds to a spatial bin defined by visual space coordinates (azimuth, elevation). For each position, a single representative receptive field is shown.

**Extended Data Fig. 4.**
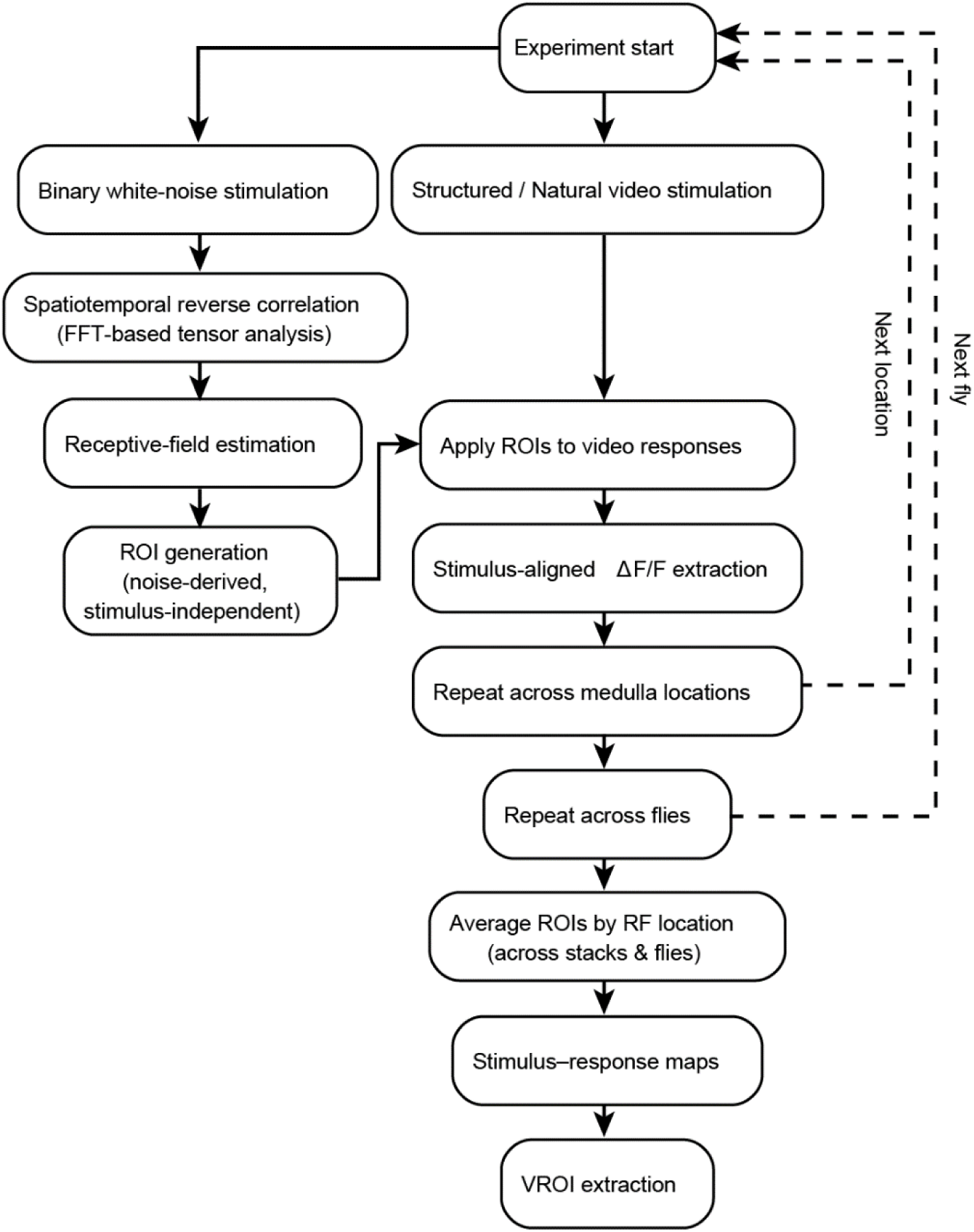
Experimental workflow for calcium imaging and data analysis. Experiments started with binary white-noise stimulation used for receptive-field estimation through spatiotemporal reverse correlation (see Methods). Receptive fields (RFs) were computed on a per-pixel basis and used to define regions of interest (ROIs) by grouping pixels with identical receptive field locations. In a separate step, structured and naturalistic videos were presented. Previously defined ROIs were used to extract stimulus-aligned fluorescence signals (*ΔF/F*) without temporal averaging over repetitions. Recordings were repeated across multiple anatomical imaging locations in each fly and across animals. Data were pooled based on receptive field location in visual space, enabling averaging of ROIs across imaging stacks and animals. The resulting population responses were used to construct stimulus–response maps in visual coordinates, which formed the basis for the extraction of virtual regions of interest (VROIs) and subsequent analyses.

**Extended Data Fig. 5.**
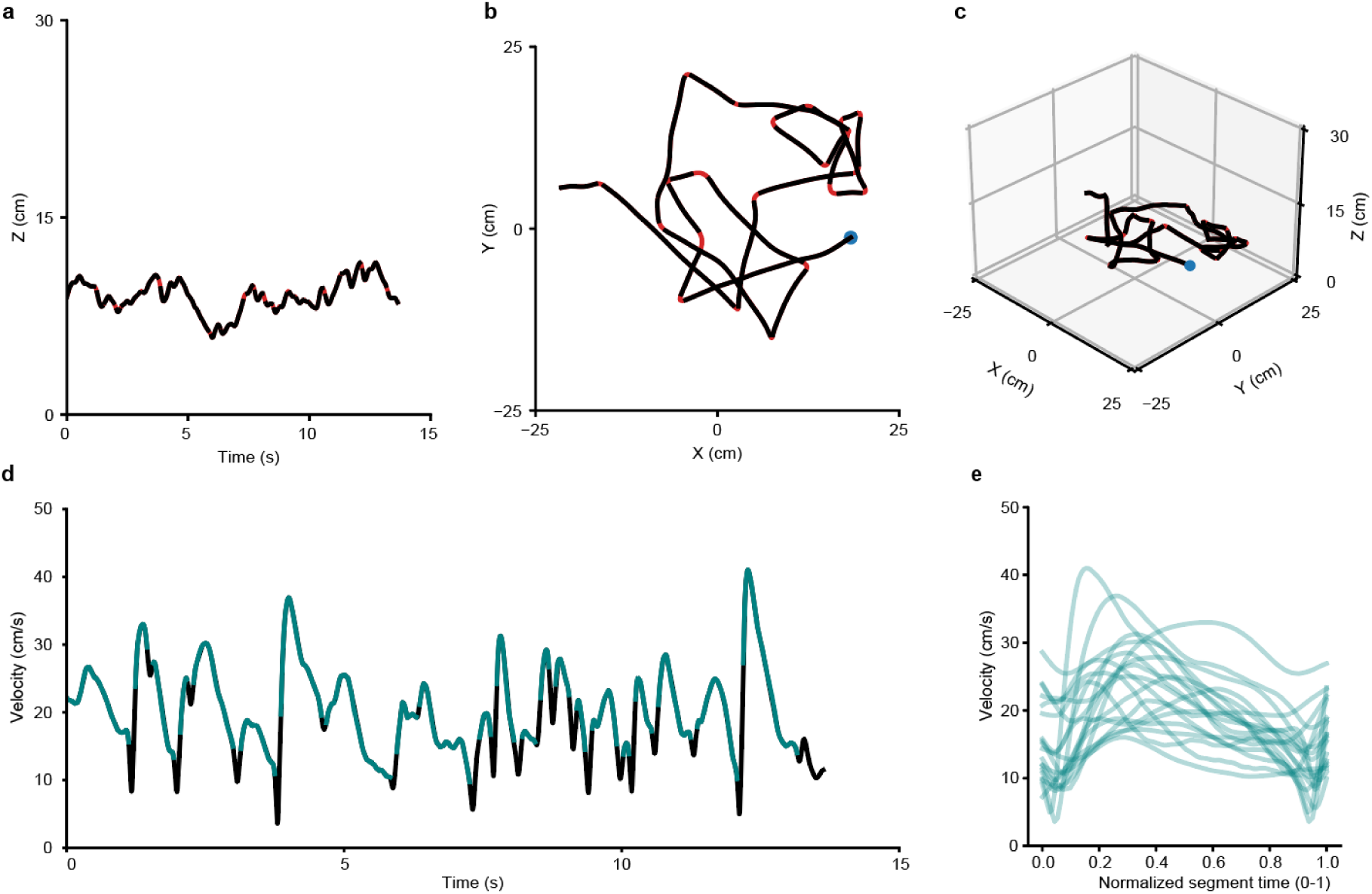
Representative free-flight trajectory. **a**, Vertical position (Z) over time for the representative trajectory. Periods of constant heading (straight flight segments) are shown in black, whereas segments with rotational changes in heading are shown in red. **b**, Two-dimensional (X–Y) trajectory of the same flight. The first tracked position is indicated in blue. Straight segments are shown in black; segments with rotational components are marked in red. **c**, Three-dimensional reconstruction of the flight trajectory (X–Y–Z). The starting position is indicated in blue. Straight segments are shown in black; segments with rotational components are marked in red. **d**, Instantaneous flight velocity over time. Teal, straight flight segments. **e**, Normalised speed profiles of straight flight segments. Individual segments are temporally rescaled to a common duration (0–1) revealing consistent velocity dynamics across segments.

**Extended Data Fig. 6.**
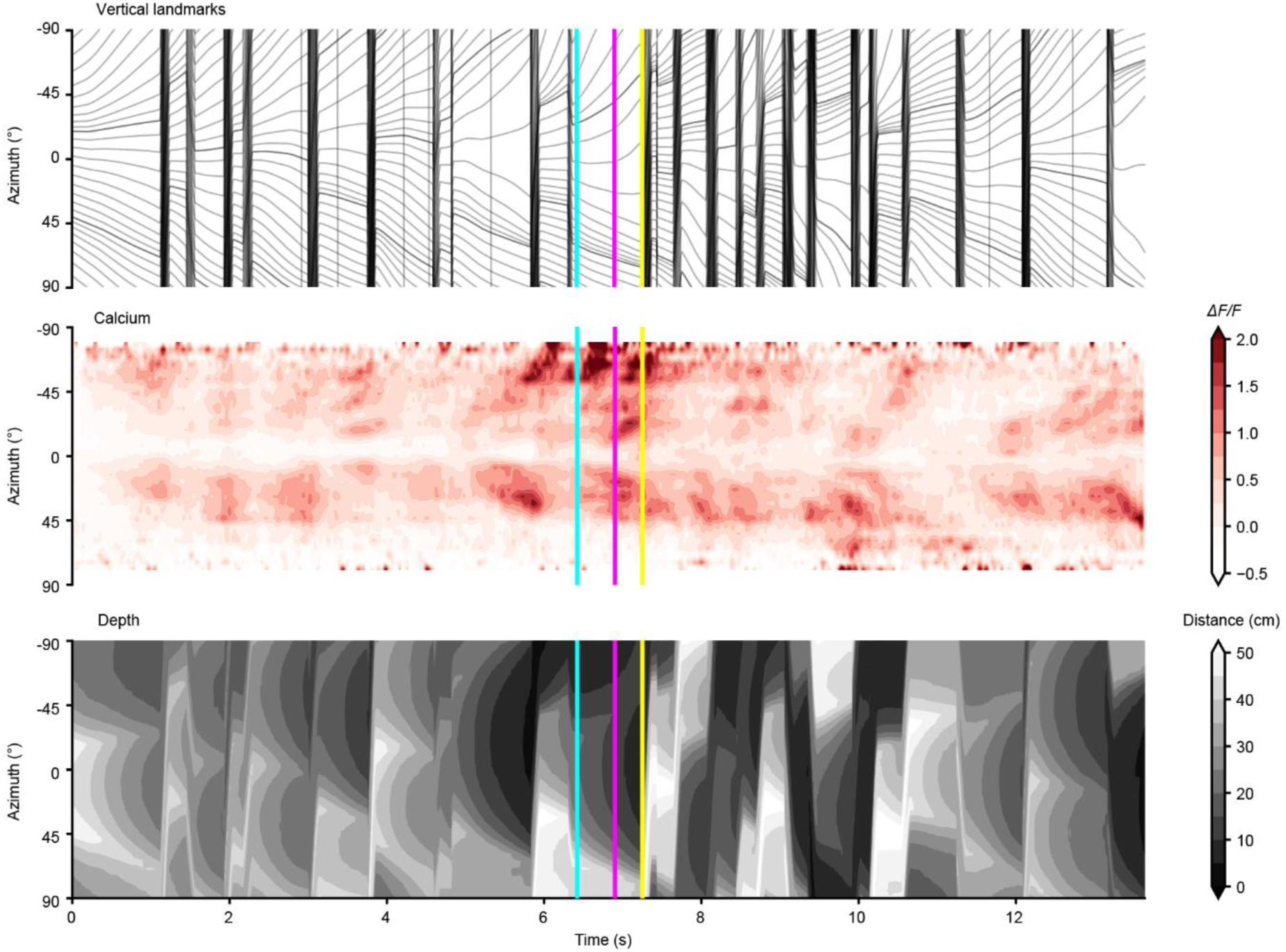
Spatio-temporal evolution of T4a activity across both hemifields. Top, reconstructed azimuthal positions of vertical landmarks over time. Each line represents the projected trajectory of an individual landmark in the visual field during virtual passage of the recorded free-flight trajectory. Centre, population-averaged calcium activity (*ΔF/F*) as a function of azimuth and time, computed by binning signals into 5° azimuthal bins and averaging across regions of interest within a defined vertical field. Bottom, corresponding depth representation of the visual scene, reconstructed from binocular stimulus videos and mapped onto the same azimuth–time coordinates. Pixel intensities represent distance (cm), averaged within the same spatial bins as the calcium signal. Vertical coloured lines (cyan, magenta, yellow) indicate example time points used for detailed analysis in Fig. 4b.

**Extended Data Fig. 7.**
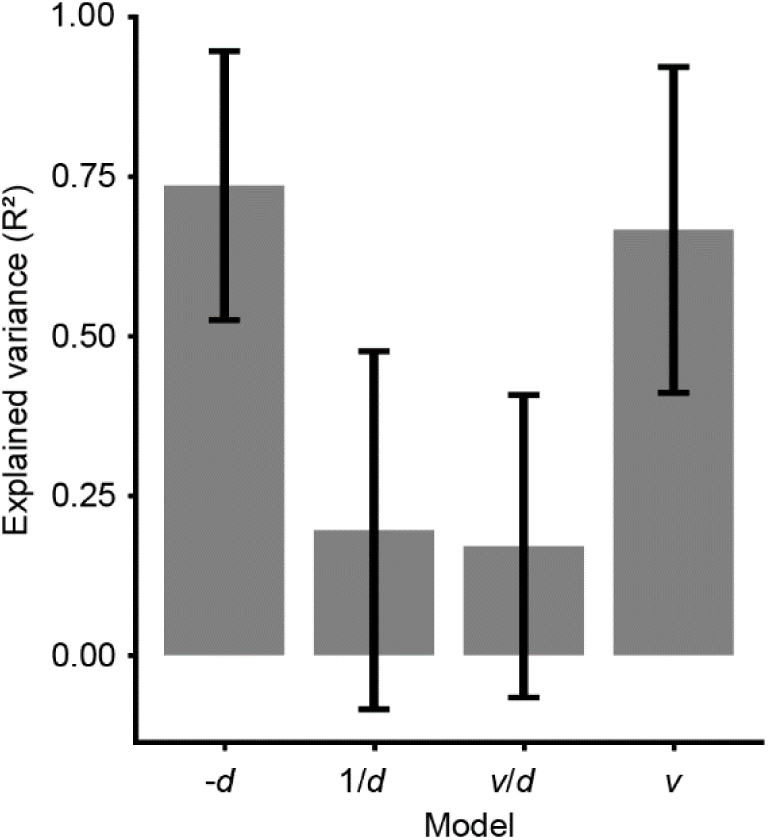
Comparison of linear models. Comparison of linear models predicting neural activity from behavioural variables. Bars indicate mean explained variance (R² ± s.d.) across segments for different predictors: distance (−*d*), inverse distance (1/*d*), optic-flow-related term (*v/d*), and velocity (*v*).

**Extended Data Fig. 8.**
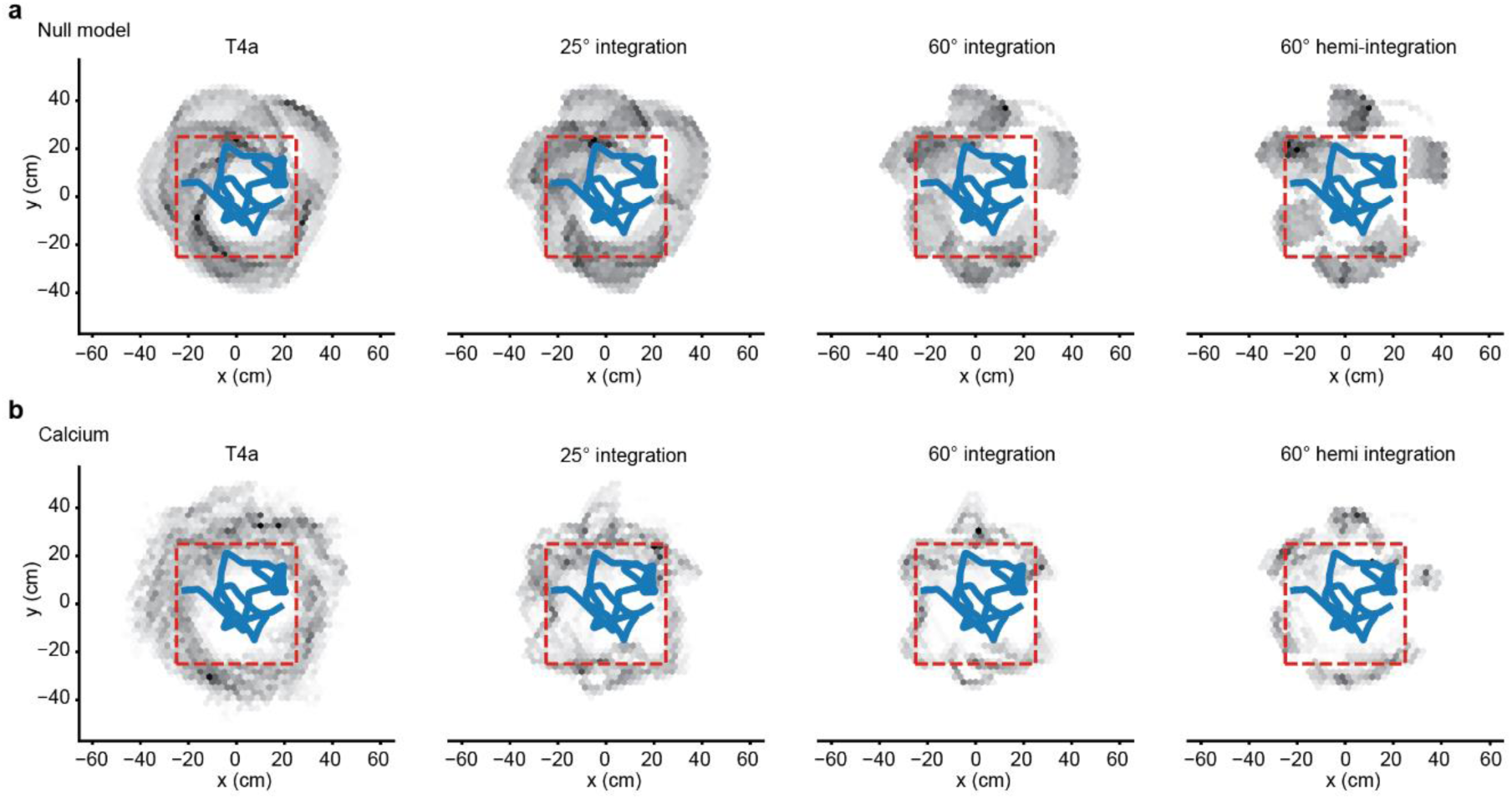
T4a calcium activity contains sufficient information to create a depth map. **a**, Null model reconstructions obtained using a constant (mean) signal without spatial information, shown for the algorithmic T4a motion detector and increasing extents of spatial integration (25°, 60°, and 60° hemispheric integration). The null model produced an isotropic, circular structure. **b**, Spatial reconstructions based on T4a calcium signals using the same integration paradigms as in **a** reveal structured approximations of the environment. Grey hexagonal bins, density of reconstructed points; blue trace, flight trajectory; dashed red square, arena boundary.

## Notes

https://github.com/stefanprech/Motion-based-depth-estimation-in-Drosophila

### Competing Interest Statement

The authors have declared no competing interest.

